# Key role of TLR3 in type I IFN expression and apoptosis induction in IBDV-infected chicken fibroblast cells

**DOI:** 10.1101/2025.07.16.665101

**Authors:** Elisabet Díaz-Beneitez, Leyre Concostrina-Martínez, Altea Martín-Martínez, Juan R. Rodríguez, José F. Rodríguez, Fernando Almazán, Dolores Rodríguez

## Abstract

Infectious Bursal Disease Virus (IBDV) (*Avibirnavirus* genus, *Birnaviridae* family) is a non-enveloped virus with a double-stranded RNA (dsRNA) genome. IBDV causes a highly contagious and immunosuppressive disease in domestic chickens (*Gallus gallus*), representing a major threat to the global poultry industry. Apoptotic cell death and exacerbated innate immune responses have been implicated in IBDV pathogenesis. Previous studies from our laboratory demonstrated the crucial role of type I interferon (IFN) in triggering apoptosis in IBDV-infected cell cultures. Genomic IBDV dsRNA is recognized by the cytoplasmic pattern recognition receptor (PRR) melanoma differentiation-associated gene 5 (MDA5) in chicken cells, triggering type I IFN responses. However, the contribution of the endosomal PRR Toll-like receptor 3 (TLR3) dsRNA sensor on type I IFN production upon IBDV infection has not been studied, despite several studies have demonstrated that its expression is significantly upregulated upon IBDV infection. Here, we demonstrate that ablation of TLR3 gene expression in DF-1 chicken fibroblasts results in a complete blockade of IBDV-induced apoptosis, a marked reduction in IFN production, and a significant enhancement of virus progeny yields. Notably, this effect appears to be specific to IBDV, as similar outcomes were not observed with other RNA viruses. Our findings also suggest that TLR3 may also play a role in viral release into the extracellular space. Additionally, receptor interacting protein kinase 1 (RIPK1), a protein that interacts with TLR3 through the adaptor Toll/IL-1 receptor (TIR) domain-containing adaptor-inducing interferon-β (TRIF), was shown to contribute to both IFN production and apoptosis in response to IBDV infection or dsRNA stimulation in DF-1 cells. Overall, this study provides new insights into the innate immune recognition of IBDV, highlighting the central role of TLR3 in mediating antiviral responses in chicken cells.

**Author summary:** Infectious Bursal Disease Virus (IBDV) is the etiological agent of a devastating syndrome known as IBD or Gumboro disease, which affects domestic chickens (*Gallus gallus*), leading to significant economic losses in the poultry industry worldwide. The virus primarily targets immature B lymphocytes causing their killing by apoptosis, resulting in severe immunosuppression. Diverse studies have suggested that exacerbation of the innate immune response is related to IBDV pathogenesis. Here, we have investigated the role of the cellular sensor of double-stranded RNA (dsRNA) Toll-like receptor 3 (TLR3) on the fate of IBDV-infected cells. Deletion of the TLR3 gene completely blocks apoptosis induced in IBDV-infected DF-1 cells, drastically reduces interferon (IFN) production and improves viral replication efficiency. We also demonstrated the participation of receptor interacting protein kinase 1 (RIPK1), a known downstream mediator of TLR3, in both, IFN production and apoptosis induction in response to IBDV infection. These findings provide new molecular insights into the mechanisms underlying the robust type I IFN response observed during IBDV infection and contribute to a deeper understanding of IBDV pathogenesis.

## Introduction

Infectious Bursal Disease Virus (IBDV), a member of the family *Birnaviridae* and the sole representative of the genus *Avibirnavirus*, is a non-enveloped virus with a bipartite double-stranded RNA (dsRNA) genome. IBDV causes a highly contagious and devastating immunosuppressive disease in young chickens (*Gallus gallus*) between 3 and 6 weeks of age. The main target cells of IBDV are immature B lymphocytes located in the bursa of Fabricius, the main lymphoid organ in birds. IBDV causes a massive depletion of infected pre-B lymphocytes and the atrophy of the bursa of Fabricius, resulting in a severe immunosuppression that predisposes chicks to secondary infections and limits the efficacy of vaccines against other important poultry pathogens. Currently, the prevalence of very virulent IBDV strains, along with the emergence of novel reassortant and recombinant strains in the field across several countries, poses a major threat to poultry industry and is responsible of significant economic losses worldwide [1–3]. Therefore, a comprehensive understanding of the mechanisms underlying IBDV-host interactions is essential for the development of effective and innovative control strategies.

IBDV virions are naked icosahedral particles containing two segments of dsRNA of 3.2 and 2.8 kbp (segments A and B, respectively). It has been proposed that upon receptor recognition, IBDV virions enter the cell through a macropinocytosis mechanism and subsequently traffic through the endocytic pathway [4,5]. During virus infection five mature viral proteins (VP1 to VP5) are synthesized. Four of these proteins are encoded by segment A, which carries two partially overlapping open reading frames (ORF). The first one (ORF A1) encodes the nonstructural protein VP5, which is dispensable for viral replication in cultured cells [6] but is essential for cell-to-cell transmission *in vitro* [7,8] and for viral pathogenesis *in vivo* [9]. The second ORF (ORF A2) encodes a polyprotein precursor, which is self-cleaved co-translationally by the viral protease VP4 [10], resulting in the formation of the VP2 precursor (pVP2), VP3 and VP4. pVP2 undergoes further cleavages to generate the mature VP2 capsid protein [11]. Proteolytic processing of pVP2 also produces four small amphipathic peptides that remain associated with the viral capsid and are believed to play a crucial role during the early stages of infection. One of these peptides, pep46, released during capsid disassembly, a process triggered by the low calcium concentration and acidic pH characteristic of mature endosomal compartments, is thought to induce pore formation in the endosomal membrane, enabling the release of the viral genome into the cytoplasm [12]. Segment B contains a single ORF that encodes VP1, a multifunctional polypeptide with an RNA-dependent RNA polymerase (RdRp) activity [13]. Within the capsid, the viral genome is structured in ribonucleoprotein complexes (RNPs), where the dsRNA is coated by VP3 and complexed with VP1 [14].

Although the molecular mechanisms underlying IBDV pathogenicity remain poorly understood, accumulating evidence indicates that the exacerbation of the innate immune response and the apoptosis of infected cells are key contributors to disease severity [15]. Significantly, previous studies from our laboratory showed that exposure to type I interferon (IFN) of HeLa cells infected with IBDV triggers a rapid and massive apoptotic cell death response, suggesting that IFN may play a critical role in the pathogenesis associated with this virus [16].

Type I IFNs are central to the host defense against viral infections. Upon binding to their membrane-bound receptor complex, IFN-Alpha/Beta Receptor (IFNAR), they initiate the Janus kinase-signal transducer and activator of transcription protein (JAK/STAT) signaling pathway, leading to the transcription of a broad repertoire of IFN-stimulated genes (ISGs). These ISGs encode proteins with diverse functions, including antiviral, immunomodulatory, cell cycle-inhibitory, and pro-apoptotic activities. Among the best-characterized pro-apoptotic ISGs is the dsRNA-activated protein kinase (PKR), which plays a key role in linking antiviral signaling to programmed cell death [17,18].

During viral infection, host cells rapidly recognize pathogen-associated molecular patterns (PAMPs), such as viral RNA or DNA, through pattern recognition receptors (PRRs), including retinoic acid-inducible gene I-like (RIG-I) receptors (RLRs) and Toll-like receptors (TLRs). Upon ligand binding, these sensors activate downstream adaptor proteins, such as the mitochondrial antiviral-signaling protein (MAVS) for RLRs, and the Toll/IL-1 receptor (TIR) domain-containing adaptor-inducing interferon-β (TRIF) or myeloid differentiation primary response 88 (MyD88) for TLRs. They initiate signaling cascades that converge on the activation of transcription factors such as IFN regulatory factors 3 and 7 (IRF3/7) and nuclear factor kappa-light-chain-enhancer of activated B cells (NF-κB), leading to the transcription of type I IFNs (IFN-α/β) and other immune effector genes [19]. In chickens, which lack the *RIG-I* gene, cytoplasmic recognition of viral dsRNA relies on melanoma differentiation-associated gene 5 (MDA5), the only RLR of this class expressed in these birds. Previous studies have shown that IBDV genomic dsRNA is sensed by MDA5, which signals through MAVS to activate IRF7—the only IRF of this type present in chickens—and NF-κB, ultimately promoting the expression of type I IFN and other proinflammatory cytokines [20–23].

As mentioned above, PKR is an ISG, but it is also a sensor for dsRNA. Our earlier work revealed that PKR can also bind IBDV dsRNA in the cytoplasm of infected cells, leading to upregulation of IFN-β expression and acting as a key mediator of the apoptotic response in IBDV infected human HeLa cells [16]. However, the role of PKR in the context of IBDV-infected chicken cells remains almost unexplored. Nonetheless, it has been shown that the IBDV-encoded VP3 protein is capable of antagonizing PKR-mediated apoptotic signaling, suggesting a potential viral strategy to modulate this pathway in chicken cells [24].

Viral dsRNA can also be recognized by the membrane associated PRRs of the TLR family. In chickens, TLR3 and TLR7 are the only TLRs known to participate in the recognition of RNA viruses, detecting dsRNA and single-stranded RNA (ssRNA), respectively [25]. In this regard, many studies have reported that the expression of the endosomal TLR3 is upregulated in different tissues of IBDV-infected chickens, as well as in infected cultured cells [26–30]. Moreover, differential regulation of TLR3 expression in the bursa, spleen and gut-associated lymphoid tissue (GALT) of chickens infected with IBDV strains of different virulence has also been reported, with higher TLR3 expression in individuals infected with more virulent strains [26,28,30–33]. Collectively, these findings suggest that the TLR3 signaling pathway plays a major role in the host immune response to IBDV and in the pathogenicity of IBDV infection. However, a direct link between TLR3 and the establishment of antiviral response against IBDV has not been reported as yet.

TLRs are type I transmembrane proteins comprising three primary domains: (i) an extracellular N-terminal ligand binding domain; (ii) a single-pass transmembrane helix that mediates membrane anchoring; and (iii) a cytoplasmic TIR domain, which facilitates protein–protein interactions essential for downstream signal transduction (Akira et al., 2006). In mammalian cells, the interaction of TLR3 with dsRNA triggers a complex signaling pathway that begins with TLR3 dimerization, the phosphorylation of a tyrosine residue in the TIR domain, and the recruitment of the adaptor protein TRIF [34].

Subsequently, TRIF facilitates the engagement of tumor necrosis factor (TNF) receptor-associated factor 3 (TRAF3), which leads to the association of two kinases, TANK binding kinase 1 (TBK1) and IkappaB kinase-epsilon (IKKε) [35] through a signalosome complex, and the activation of IRF3/7, while the recruitment of TRAF6 and the receptor interacting protein kinase 1 (RIPK1) triggers the activation of NF-kB [36]. In chicken cells, the signaling pathway downstream of TLR3 remains poorly characterized. Nonetheless, the identification of functional homologs for most key mammalian components, including TRIF, TBK1, IKKε, and RIPK1, in the chicken genome (*Gallus gallus*) suggests a conserved signaling mechanism across these species [37].

In this study, we investigated the molecular mechanisms underlying apoptosis induction in IBDV-infected chicken cells. Our findings reveal a strong link between IFN-mediated activation of the JAK/STAT signaling pathway and the initiation of apoptosis. We also examined the contribution of the cytoplasmic and endosomal dsRNA sensors—MDA5, PKR, and TLR3—to IFN expression and cell death. Our results demonstrate that the three PRRs participate in this process; however, TLR3 emerges as the primary mediator, as its ablation completely abrogates apoptosis in IBDV-infected cells. Furthermore, we have identified the downstream signaling molecule RIPK1 as a key effector in TLR3-mediated apoptotic signaling.

## RESULTS

### Interplay between IFN production and apoptosis induction during IBDV infection

Previous research from our laboratory demonstrated that type I IFN plays a pivotal role in the fate of IBDV-infected cells. Specifically, while pretreatment of cells with IFN-α provides a robust protection against IBDV replication, its addition early after infection triggers a significant apoptotic response that effectively eliminates infected cell cultures [16]. Moreover, recent research has demonstrated that the inactivation of the JAK/STAT pathway markedly attenuates IBDV-induced cell death in avian DF-1 cells, which in turn, facilitates the establishment of persistent IBDV infections [38].

In this study, we employed ruxolitinib (Rx), a selective inhibitor of the JAK/STAT signaling pathway, to assess the contribution of endogenous type I IFN to the apoptotic response observed in IBDV-infected DF-1 cells that result in the massive cell death observed late in infection. To this end, mock-infected or IBDV-infected DF-1 cells were treated 1 hour (h) after virus adsorption with Rx at a concentration of 4 μM. All infections throughout this study were performed at a multiplicity of infection (MOI) of 2 plaque forming units (PFU) per cell (PFU/cell), and cells were collected at two time points, 16 and 24 h post-infection (pi) for subsequent analyses. In parallel, additional sets of mock-infected and IBDV-infected cultures were treated with 1,000 U/mL of recombinant chicken IFN-α (chIFN-α) at 3 h pi and harvested at 16 h pi. As previously reported, exogenous IFN treatment enhanced the cytopathic effect in IBDV-infected cells at 16 h pi, yielding a phenotype comparable to that of untreated infected cells at 24 h pi (data not shown). To confirm that this effect corresponded to apoptotic cell death, the activity of caspases 3 and 7 in the cultures was analyzed with the commercial Caspase-Glo 3/7 kit (Fig 1). The results demonstrated that the levels of apoptosis observed in cells infected for 24 h were comparable to those observed in IFN-treated infected cells collected at 16 h pi. (Fig 1A). In both cases, apoptosis was significantly diminished in Rx-treated cell cultures. Then, the expression of the *IFNB* gene in these samples was analyzed by real-time quantitative PCR (RT-qPCR) to explore the relationship between IFN signaling and apoptosis. As shown in Fig 1B, high levels of induction of *IFNB* were observed in infected DF-1 cells harvested at 24 h pi, being comparable to those detected in IFN-treated infected cells harvested at 16 h pi. However, in all infected cells treated with Rx there was a notable reduction in the expression of this gene (ca. 3 log units). Similar expression patterns were observed for the *MDA5*, *TLR3* and *Mx* ISGs (Figs. 1C-E). These data, indicate the existence of a feedback mechanism whereby IFN-β, secreted by infected cells, stimulates the expression of the gene itself as well as those of the ISGs. The use of Rx confirms that inhibiting the type I IFN signaling pathway results in a significant reduction in apoptosis in IBDV-infected cells, suggesting that IFN released by IBDV-infected cells is responsible for the apoptosis observed at late times pi.

**Fig 1.**
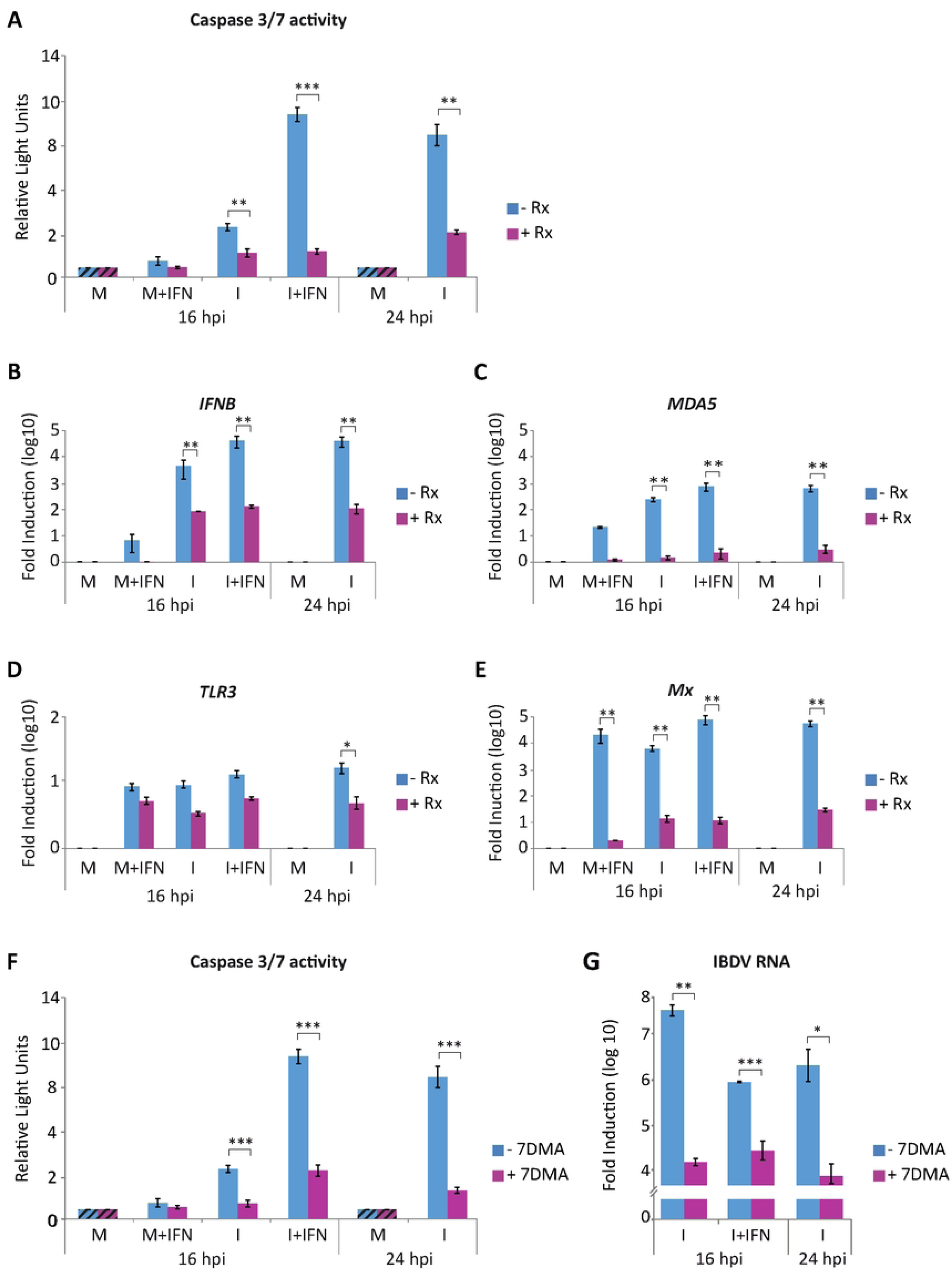
Treatment with either the JAK/STAT inhibitor ruxolitinib (Rx) or the viral polymerase inhibitor 7DMA significant decrease apoptotic death of IBDV-infected DF-1 cells. DF-1 cells mock-infected (M) or infected with IBDV (MOI of 2 PFU/cell) (I) were treated with Rx (4 µM) (A-E) or with 7DMA (0.2 mM) (F-G) after virus adsorption, then treated or not with chIFN-α (1,000 IU/ml) at 3 h pi (M+IFN, I+IFN), and collected at 16 and 24 h pi. (A) Apoptosis was measured by using the Caspase-Glo 3/7 assay kit. Each determination was carried out in duplicate. Caspase values from infected cell samples were normalized to those from mock-infected cells (M) (striped bars). (B-E) The expression levels of *IFNB* (B), *MDA5* (C), *TLR3* (D) and *Mx* (E) genes were determined by SYBR green-based RT-qPCR. Recorded values for each cellular gene were normalized to the *GAPDH* mRNA content and presented on a log_10_ scale as the fold induction over the level found in mock-infected DF-1 cells. (F) Apoptosis was measured by using the Caspase-Glo 3/7 assay kit, and as above, caspase values from infected cell samples were normalized to those from mock-infected cells (M) (striped bars). (G) The accumulation levels of the IBDV RNA were determined by SYBR green-based RT-qPCR. Bars indicate means ± standard deviations based on data of duplicate samples from three independent experiments. *, ** and *** indicate *p* values of <0.05, <0.01 and <0.001 respectively, as determined by unpaired Student’s test.

### Apoptosis of IBDV-infected cells is dependent upon viral replication

Previous results obtained in HeLa cells indicated that viral RNA replication/transcription is required for massive apoptosis induction upon IFN treatment of IBDV-infected human HeLa cells [16]. Then, to confirm that viral RNA replication/transcription is also required in the context of IBDV-infected chicken cells we used the viral RNA polymerase inhibitor 7-deaza-2’-C-metiladenosina (7DMA). For this, mock-infected or cells infected with IBDV were treated 1 h after virus-adsorption with 7DMA at a concentration of 0.2 mM and collected at 16 and 24 h pi for analysis. As above, a portion of the cultures collected at 16 h pi had been treated with chIFN-α (1,000 U/ml) at 3 h pi. A drastic decrease in caspase 3/7 activity was detected in samples from 7DMA-treated cells, both in those from cultures just infected and harvested at 16 and 24 h pi, as well as in samples from IFN-treated cell cultures, compared to cells not treated with 7DMA (Fig 1F). The effectiveness of the 7DMA treatment was assessed by the analysis of viral RNA levels by RT-qPCR. As shown in Fig 1G, an overall reduction of ca. 2 to 3 log units was determined in all samples treated with 7DMA. These results indicate that the replication/transcription process of viral dsRNA is a critical determinant for apoptosis induction also in IBDV-infected DF-1 cells.

### MDA5 and MAVS contribute to the activation of apoptosis in IBDV-infected DF-1 cells

As mentioned above, results from different laboratories, including ours, showed that in chicken cells IBDV dsRNA is recognized by the cytoplasmic sensor MDA5 leading to the recruitment of the adaptor protein MAVS and the activation of the IFN signaling pathway. We therefore set out to investigate the possible involvement of MDA5 and MAVS proteins in the apoptotic process triggered during IBDV infection by using the RNA interference-mediated gene silencing technology. After optimizing transfection conditions (data not shown), DF-1 cells were transfected with specific small interfering RNAs (siRNAs) targeting *MDA5* and *MAVS* genes, either independently or simultaneously, as described in the Materials and Methods section. Under optimal conditions, we achieved a 75-90% reduction in the mRNA expression of both genes compared to cells transfected with a control siRNA (Fig 2A).

**Fig 2.**
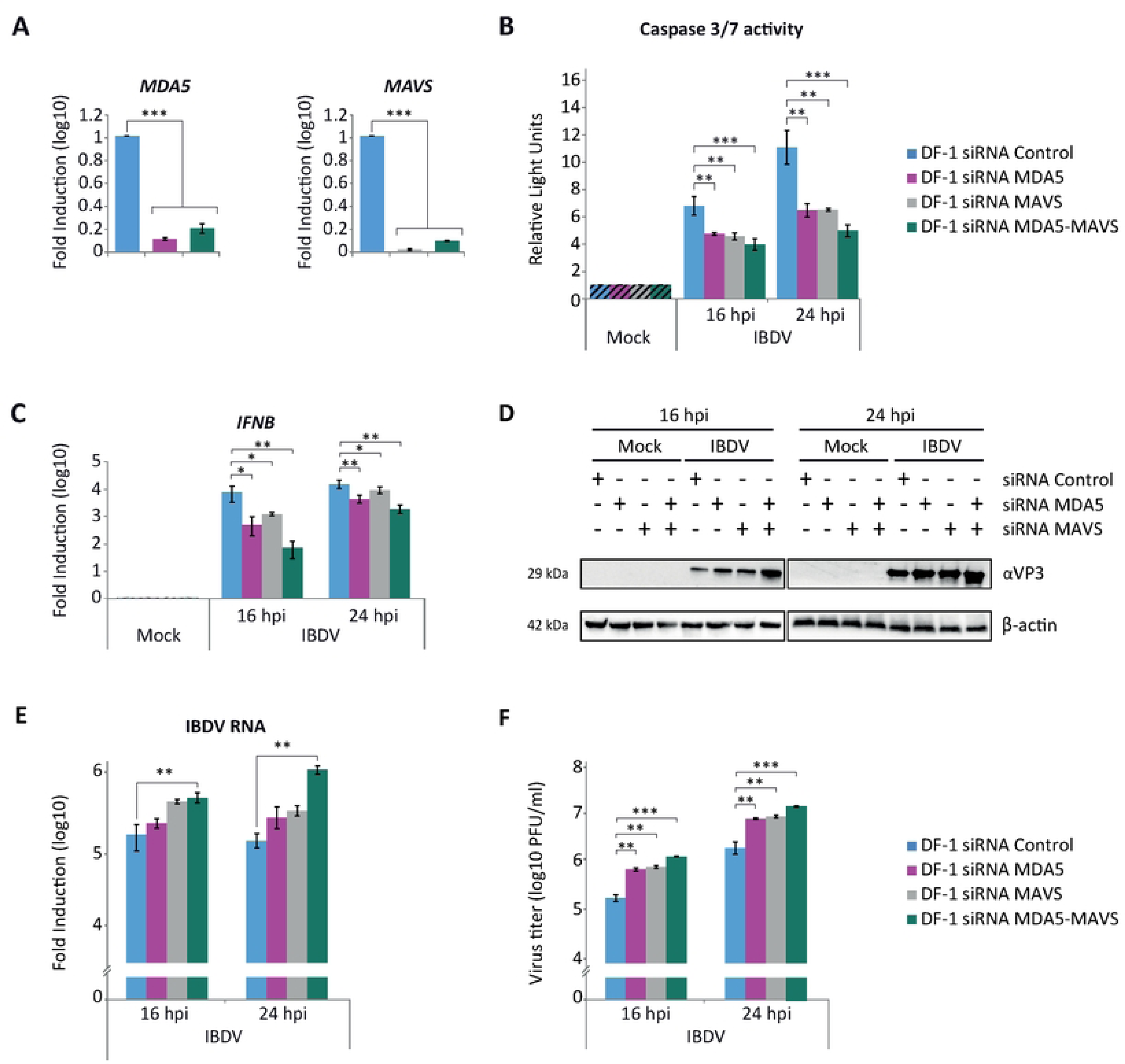
MDA5 and MAVS contribute to induce apoptosis in DF-1 cells infected with IBDV. DF-1 cells were transfected with siRNAs that silence the expression of MDA5 and MAVS proteins or with a control siRNA. At 24 h pt, cell cultures were infected or not (Mock) with IBDV (MOI 2 PFU/cell), and cell samples were collected at 16 and 24 h pi. **(A)** Extracts from uninfected cells collected at 24 h pt were used for total RNA extraction and subsequent analysis of *MDA5* and *MAVS* mRNA levels by RT-qPCR using specific primers for the silenced genes. The expression values of the *MDA5* and *MAVS* genes were normalized to the *GAPDH* gene. Values were relative to those obtained in cultures transfected with the control siRNA. **(B)** Apoptosis was measured by using the Caspase-Glo 3/7 assay kit, and each determination was carried out in duplicate. Caspase values from infected cell samples were normalized to those from mock-infected cells (Mock) (striped bars). **(C)** RT-qPCR study of the expression of *IFNB* gene. Cellular gene expression values were normalized to those of the *GAPDH* mRNA content and are presented on a log_10_ scale as the fold induction over the level found in mock-infected DF-1 silenced control cells. **(D)** Immunoblot analysis of total cell extracts with antibodies specifically recognizing the IBDV structural VP3 protein. Antibodies to β-actin were used for protein loading control. **(E)** The accumulation levels of the IBDV RNA were determined by SYBR green-based RT-qPCR. **(F)** Extracellular virus yields. Bars indicate means ± standard deviations based on data of duplicate samples from three independent experiments. *, ** and *** indicate *p* values of <0.05, <0.01 and <0.001, respectively, as determined by unpaired Student’s test.

At 24 h post-transfection (pt), silenced DF-1 cells were infected with IBDV and harvested at 16 and 24 h pi. Phase-contrast microscopy revealed a marked reduction in the cytopathic effect (CPE) in MDA5- or MAVS-silenced cells compared to control cells at both time points (data not shown). This observation was corroborated by the analysis of caspases 3/7 activation, which revealed a reduction in apoptosis of approximately 1.5-fold in cells silenced with either *MDA5* or *MAVS* siRNAs at 24 h pi. Notably, co-silencing of both genes resulted in a reduction of ca. 2-fold compared to cells silenced with the control siRNA (Fig 2B). Consistently, RT-qPCR analysis of *IFNB* gene expression showed significantly reduced transcript levels in all siRNA-treated infected cultures, with the strongest suppression observed in the double-silenced condition (Fig. 2C).

Regarding IBDV replication, a notable increase in the accumulation of the viral VP3 protein was observed in DF-1 cells transfected with *MDA5* or *MAVS* siRNAs, with a further enhancement in cells transfected with both siRNAs (Fig 2D). These results were then complemented with the quantification of viral RNA accumulation by RT-qPCR and the determination of viral titers in cell supernatants. Both the accumulation of IBDV RNA in cells and the titers of the virus in the culture supernatants were found to be significantly higher in samples from silenced cells with either *MDA5* or *MAVS* siRNA, at both 16 and 24 h pi, compared to cells infected after transfection with control siRNA. Again, this difference was accentuated when both siRNAs were used simultaneously (Figs. 2E-F).

Collectively, these results indicate that silencing *MDA5* and *MAVS* genes in DF-1 cells prior to IBDV infection results in a significant, but partial reduction in the activation of apoptosis, a decrease in the activation of the innate immune response and an increase in the efficiency of viral replication.

### PKR contributes significantly to apoptosis induction in IBDV-infected DF-1 cells

PKR plays a pivotal role in cellular responses to diverse stress-inducing stimuli, including viral infections, and is involved in apoptosis induction through various mechanisms [17,18]. In this regard, we previously showed that PKR acts as a critical mediator of the apoptotic response in IBDV-infected human HeLa cells treated with IFN-α [16]. Then, we sought to study the potential contribution of PKR to apoptosis triggering in IBDV-infected chicken DF-1 cells. For this, we generated a DF-1 PKR KO cell line using the CRISPR/Cas9 technology as described in the Materials and Methods section. We designed two guides to produce a double cut in the target gene that should result in a large deletion entailing the loss of the sequence coding for the two N-terminal dsRNA binding domains and a portion of the active site of the protein, as depicted in Fig 3A. Four cell clones were selected, and the expression of PKR was assessed by Western blot analysis in comparison with WT DF-1 cells (Fig 3B). No expression of PKR was detected in any of the clones, while the protein present in WT cells was clearly detected. To further characterize these cells, genomic DNA was extracted and subjected to sequencing analysis, which revealed that only clone 3 presents a deletion of the expected size (10.107 Kb), while in the other three clones there had occurred rearrangements of the sequence most likely resulting from non-homologous recombination repair. DF-1 PKR KO cells from clone 3 were then used to determine the role of PKR in apoptosis induction in IBDV-infected chicken cells. For this, WT and DF-1 PKR KO cells were infected and samples were taken at 16 and 24 h pi for the analysis of apoptotic cell death by determining caspase 3/7 activity. We observed a significant decrease in the apoptotic levels in DF-1 PKR KO as compared with WT DF-1 cells, ranging from about 30% reduction at 16 h pi to 60% at 24 h pi (Fig 3C). Furthermore, by RT-qPCR we observed a concomitant reduction in *IFNB* gene expression in IBDV-infected DF-1 PKR KO cells compared with WT DF-1 cells at both time points (Fig 3D). These results show that PKR plays a major role in the induction of apoptosis upon IBDV infection, that could be directly or indirectly related with IFN-β production. However, the persistence of a significant level of apoptotic cell death in the absence of the PKR protein indicates that other factor(s) is also involved in IBDV-induced apoptosis in chicken cells.

**Fig 3.**
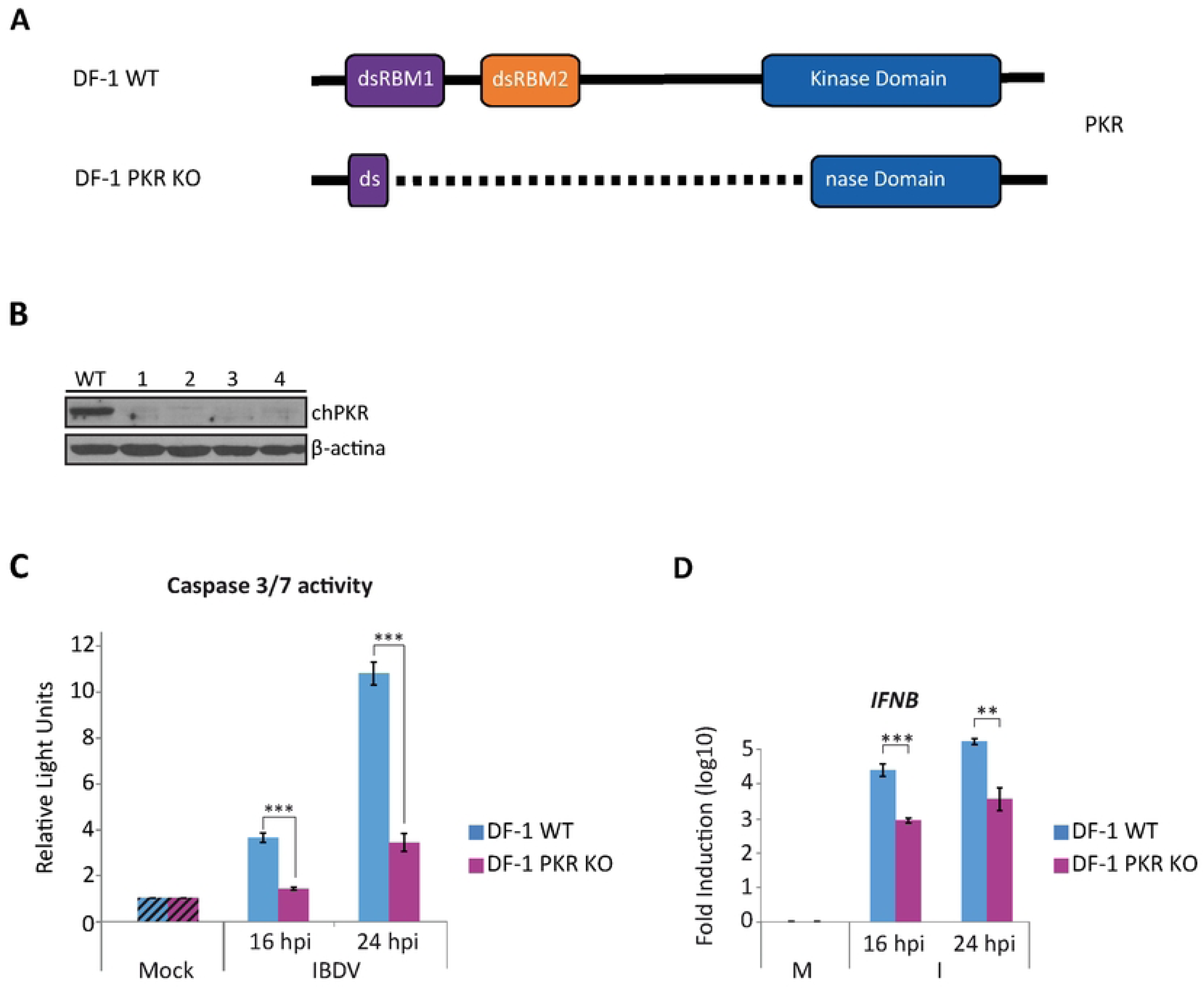
The induction of apoptosis by IFN in IBDV-infected DF-1 cells is partially dependent on PKR expression. **(A)** Schematic representation of the structure of the WT *PKR* gene and the deleted version of the gene present in the DF-1 PKR KO cells. **(B)** Western blot analysis of total cell extracts with antibodies specifically recognizing the PKR protein. Antibodies to β-actin were used for protein loading control. **(C-D)** DF-1 and DF-1 PKR KO cells were mock-infected or infected with IBDV (MOI of 2 PFU/cell) and at 16 and 24 h pi the levels of apoptosis **(C)** and the expression of *IFNB* gene **(D)** were analyzed by using the Caspase-Glo 3/7 assay kit and RT-qPCR, respectively. Caspase activity was determined in duplicate, and the values normalized to those from mock-infected cells (Mock) (Striped bars). Cellular gene expression values were normalized to those of the *GAPDH* mRNA content and are presented on a log_10_ scale as the fold induction over the level found in mock-infected DF-1 or DF-1 PKR KO cells. Bars indicate means ± standard deviations based on data of duplicate samples from three independent experiments. ** and *** indicate *p* values of <0.01 and <0.001, respectively, as determined by unpaired Student’s test.

### TLR3 is an essential factor in the induction of apoptosis in IBDV-infected DF-1 cells

As introduced earlier, TLR3 is a transmembrane receptor located mainly on endosomes, where it recognizes viral dsRNA and initiates innate immune signaling cascades. It has been proposed that IBDV enters the cell through a macropinocytosis mechanism [5] and internalizes into endosomal vesicles where the virus capsid is destabilized and RNPs are released. It is therefore reasonable to assume that the TLR3 receptor might act as the first cellular sensor capable of recognizing IBDV dsRNA upon virus entry into the cells. This, together with the observation that *TLR3* gene expression is induced in IBDV-infected DF-1 cells as well as in bursal cells from infected animals [26–30], led us to investigate the role of this transmembrane receptor during the IBDV infection process. For this, we generated TLR3-deficient DF-1 cell lines using the CRISPR/Cas9 technology, employing two paired combinations of 3 guide RNAs, that allowed us to generate two different partial deletions affecting the extracellular domain of the protein responsible for dsRNA recognition. As depicted in Fig 4A, two representative cell clones, 7 and 10, were selected. Sequencing analysis revealed that clone 7 harbors a 949 bp (nucleotides 1081-2030) deletion in the *TLR3* gene, causing frameshifts and the predicted expression of a defective protein, while clone 10 presents a deletion of 1,697 bp (nucleotides 339-2036) affecting most of the extracellular domain. Unfortunately, attempts to detect the TLR3 protein in samples from mock-infected or IBDV-infected DF-1 WT cells by Western blot using several commercial antibodies failed. Although we could not confirm the absence of the target protein in the DF-1 TLR3 KO cell clones by Western blot, the extent and nature of the corresponding gene deletions strongly suggest a complete loss of functional TLR3 expression in these CRISPR-edited cells.

**Fig 4.**
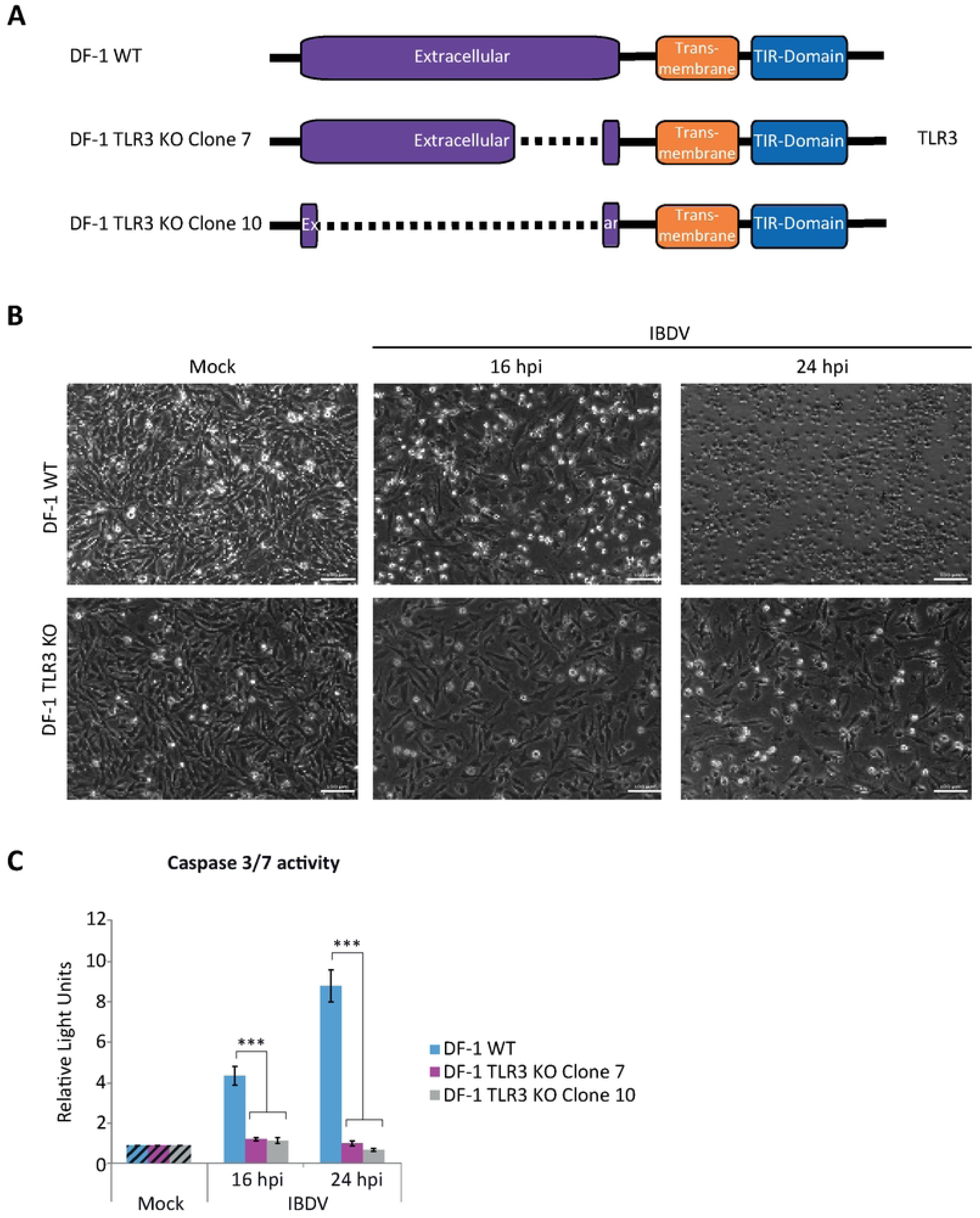
Apoptosis triggering in IBDV-infected DF-1 cells is dependent on TLR3 expression. **(A)** Schematic representation of the structure of the *TLR3* gene and the deleted versions of the gene present in two selected DF-1 TLR3 KO cell clones. **(B-C)** DF-1 and DF-1 TLR3 KO cells were mock-infected or infected with IBDV (MOI of 2 PFU/cell) and at 16 and 24 h pi the extent of CPE **(B)** and the levels of apoptosis **(C)** were analyzed by phase-contrast microscopy and using the Caspase-Glo 3/7 assay kit, respectively. Caspase activity was determined in duplicate and the values normalized to those from mock-infected cells (Mock) (Striped bars). Bars indicate means ± standard deviations based on data of duplicate samples from three independent experiments. *** indicate *p* value of <0.001, as determined by unpaired Student’s test.

Once the DF-1 TLR3 KO cell lines were generated, our objective was to ascertain the effect of the TLR3 protein ablation on the IBDV infection process. To this end, DF-1 WT and DF-1 TLR3 KO cells from both clones were infected with IBDV and samples were taken at 16 and 24 h pi for various analyses. Phase contrast microscopy revealed a significant reduction in the cytopathic effect of the infected DF-1 TLR3 KO cells, particularly at 24 h pi (Fig 4B). This observation was corroborated by the analysis of caspase 3/7 activity. As shown in Fig 4C, there was a drastic reduction in the level of apoptosis detected in infected cells from both TLR3 KO cell lines in comparison to DF-1 WT cells at both times pi. However, the impact was more pronounced at 24 h pi, when higher levels of apoptosis are detected in DF-1 WT cells. Indeed, the levels of apoptosis in the two DF-1 TLR3 KO cell lines infected with IBDV are comparable to those found in uninfected (mock) cells, as well as to those detected in WT infected cells treated with the pan-caspase inhibitor Z-VAD-FMK (S1 Fig). However, both cell lines undergo high levels of apoptosis, comparable to those of WT DF-1 cells after treatment with staurosporine, a well-known apoptotic inducer used as a control (S1 Fig). Strikingly, these findings demonstrate that TLR3 silencing alone is sufficient to fully abrogate apoptosis induced by IBDV infection in DF-1 cells, underscoring its pivotal role in orchestrating the apoptotic response.

### Role of TLR3 in the control of viral replication and dissemination

We next wanted to analyze the impact of the TLR3 deficiency on IBDV replication. Using the same infection conditions described above, we analyzed the accumulation of the viral protein VP3 by Western blot. A significant increase in VP3 accumulation was clearly observed in samples obtained from both DF-1 TLR3 KO cell lines compared to DF-1 WT cells at both time points pi (Fig 5A). To confirm these findings, we performed an RT-qPCR analysis to assess the accumulation of viral RNA. We found that in the absence of TLR3, the amount of viral RNA increased by about 1 log unit or slightly more at 16 and 24 h pi, respectively (Fig 5B). However, the analysis of the accumulation of infectious virus in the culture supernatants revealed a significant decrease in viral titers obtained in samples from the DF-1 TLR3 KO cell lines compared to the parental cells (Fig 5C). These results appear to indicate a reduction in viral replication and are therefore in apparent contradiction with the findings from the VP3 protein analysis by Western blot and the quantification of viral RNA by RT-qPCR. To try to solve this discrepancy, we quantified the intracellular virus titers in the different infected cultures. The results showed that the intracellular viral titers obtained from the DF-1 TLR3 KO cells were significantly higher than those obtained from the DF-1 WT cells at both time points pi (Fig 5D), concurring with the protein and RNA data. Taken together, this set of data demonstrate that TLR3 is a key player in the apoptotic process triggered during IBDV infection in DF-1 cells, as well as in IBDV replication. In addition, a potential role in virus release is suggested by the reduction of extracellular viral titers observed in the DF-1 TLR3 KO cells. These results are reminiscent of those obtained with an IBDV mutant virus, the VP5 KO virus, generated in our laboratory by reverse genetics that does not express the viral protein VP5. Our previous research demonstrated that the VP5 protein plays a role in the extracellular release of virions during the early stages of the viral life cycle, through a non-lytic mechanism [8].

**Fig 5.**
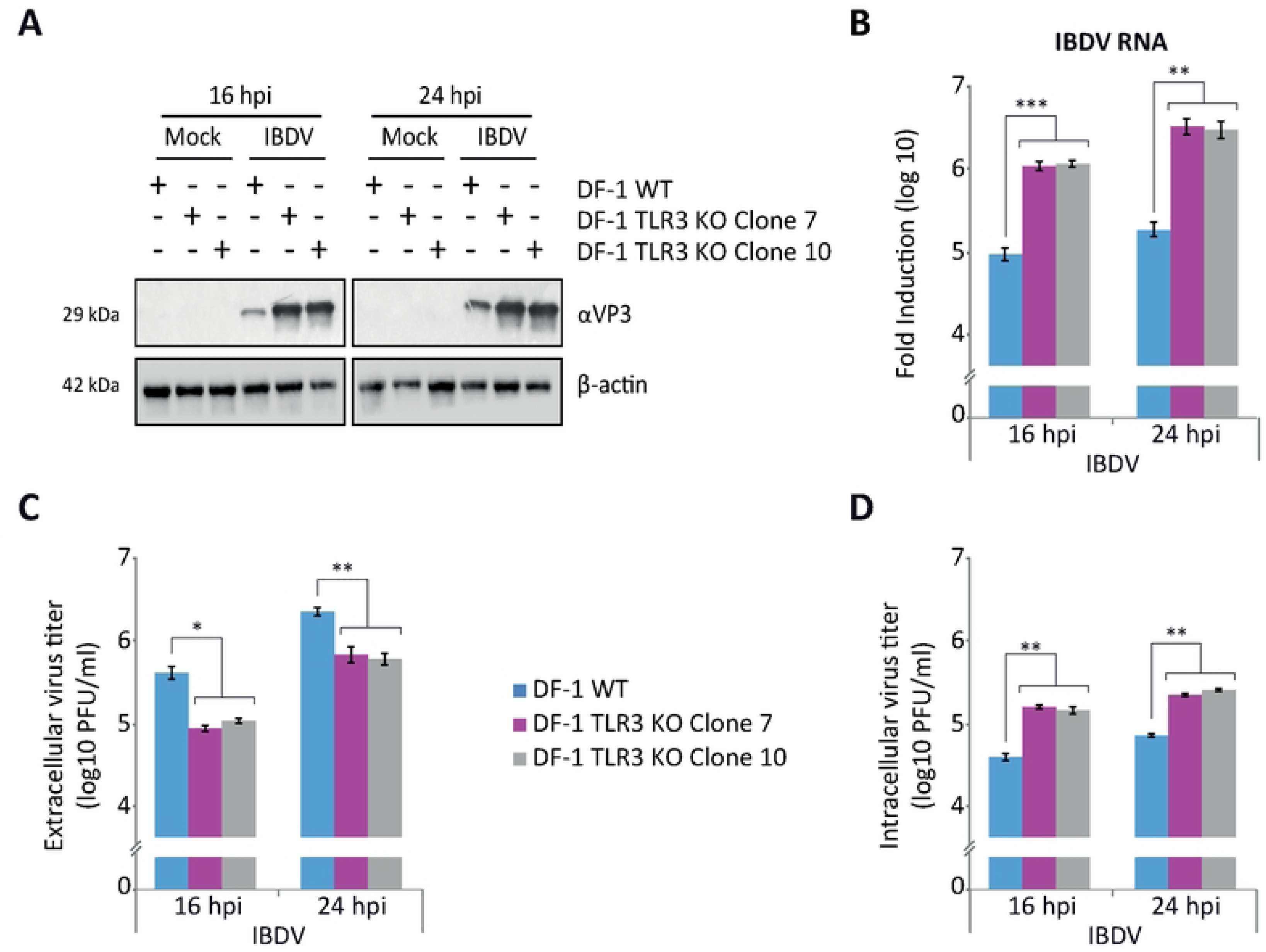
TLR3 is an essential factor in the control of IBDV replication. DF-1 and DF-1 TLR3 KO cells were mock-infected or infected with IBDV (MOI of 2 PFU/cell). Cell samples were harvested at 16 and 24 h pi. **(A)** Western blot analysis of total cell extracts with antibodies specifically recognizing the IBDV structural VP3 protein. Antibodies to β-actin were used for protein loading control. **(B)** The accumulation of the IBDV RNA was determined by SYBR green-based RT-qPCR. Analysis of extracellular **(C)** and intracellular **(D)** virus titers. Bars indicate means ± standard deviations based on data of duplicate samples from three independent experiments. *, ** and *** indicate *p* values of <0.05, <0.01 and <0.001, respectively, as determined by unpaired Student’s test.

To compare the impact of VP5 absence in the virus with the absence of TLR3 in the cells, WT DF-1 and DF-1 TLR3 KO cells were infected with both, WT and VP5 KO viruses, and samples corresponding to cell cultures and supernatants were collected at 8 and 16 h pi for quantification of viral titers. In the case of IBDV VP5 KO, no differences in viral titers were observed between supernatants from the DF-1 TLR3 KO and the DF-1 WT cell lines at either time pi (S2A Fig). However, as previously observed, at 16 h pi the titers of WT virus in the supernatants from DF-1 TLR3 KO cells were significantly lower than in those from the DF-1 WT cells, but comparable with those obtained in both cell lines infected with IBDV VP5 KO. At 8 h pi there were no significant differences between the different samples. These results indicate that the egression of IBDV WT virus is reduced in DF-1 TLR3 KO cells, as it is for IBDV VP5 KO virus in both cell lines. In contrast, when analyzing intracellular virus production, we again observed that virus titers were higher in DF-1 TLR3 KO cells infected with IBDV WT than in DF-1 WT cells at both 8 and 16 h pi (S2B Fig). In the case of the IBDV VP5 KO virus, we also observed an increase in intracellular virus production in samples from DF-1 TLR3 KO cells, although at 16 h pi the differences were not statistically significant. Overall, the results obtained in DF-1 TLR3 KO cells are consistent with those using the IBDV VP5 KO virus, suggesting that TLR3 contributes not only to the antiviral response but also to viral dissemination.

As the results obtained with both DF-1 TLR3 KO cell lines are similar, we decided to use only clone 10 for further analyses.

### The TLR3-initiated pathway is primarily responsible for IFN production in IBDV-infected DF-1 cells

The next objective was to analyze *IFNB* gene expression during IBDV infection in DF-1 TLR3 KO cells. DF-1 WT and DF-1 TLR3 KO cells were infected and total RNA was isolated at 16 and 24 h pi for RT-qPCR analysis. As shown in Fig 6A, the induction of *IFNB* was significantly lower (ca. 2 log units) in samples from DF-1 TLR3 KO cells than in DF-1 WT cells at both time points pi. To further assess the impact of TLR3 deficiency on the antiviral response, we expanded the study to analyze the expression of several ISGs, including *MDA5*, *OAS* and *Mx*. The results confirmed the reduced innate immune response during the IBDV infection process in DF-1 TLR3 KO cells at both 16 and 24 h pi compared to parental cells (Figs. 6B-6D).

**Fig 6.**
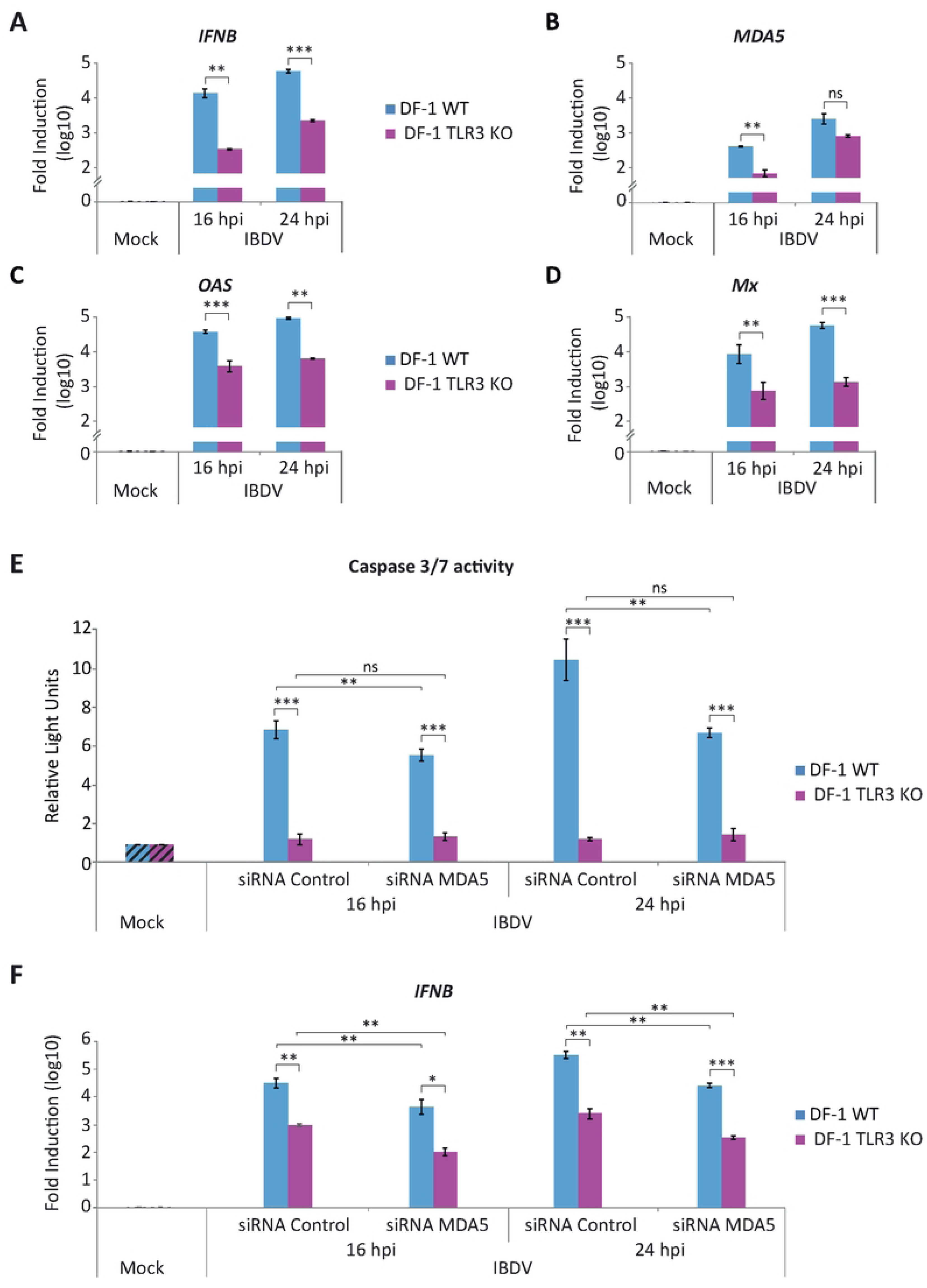
Activation of the innate immune response during IBDV infection in DF-1 cells. **(A-D)** DF-1 and DF-1 TLR3 KO cells were mock-infected or infected with IBDV (MOI of 2 PFU/cells) and cells were harvested at 16 and 24 h pi for RNA extraction and RT-qPCR analysis. The expression levels of *IFNB* **(A)**, *MDA5* **(B)**, *OAS* **(C)** and *Mx* **(D)** genes were determined by SYBR green-based RT-qPCR. Recorded values for each cellular gene were normalized to the *GAPDH* mRNA content and are presented on a log_10_ scale as the fold induction over the level found in mock-infected DF-1 or DF-1 TLR3 KO cells. **(E-F)** DF-1 cells were transfected with a siRNA specifically silencing the expression of MDA5 protein, or with a control siRNA. At 24 h pt, cells were either infected or not (Mock) with IBDV (MOI 2 PFU/cell). Samples were collected at 16 and 24 h pi and the levels of apoptosis **(E)** and the expression of the *IFNB* gene **(F)** analyzed by using the Caspase-Glo 3/7 assay kit and RT-qPCR, respectively. Caspase activity was determined in duplicate, and the values normalized to those from mock-infected cells (Mock) (Striped bars). Cellular gene expression values were normalized to those of the *GAPDH* mRNA content and are presented on a log_10_ scale as the fold induction over the level found in mock-infected DF-1 or DF-1 TLR3 KO cells. Bars indicate means ± standard deviations based on data of duplicate samples from three independent experiments. *, ** and *** indicate *p* values of <0.05, <0.01 and <0.001, respectively, as determined by unpaired Student’s test. ns, not significant.

Then, we wanted to investigate the combined effect of TLR3 and MDA5 on the activation of the innate immune response induced in IBDV-infected DF-1 cells. To achieve this, we transfected DF-1 WT and DF-1 TLR3 KO cells with a MDA5-specific siRNA. Once we had confirmed that the silencing process had worked correctly, with a reduction of approximately 70% in *MDA5* mRNA compared to cells transfected with the control siRNA (data not shown), we proceeded to infect the cells with IBDV. Samples were taken at 16 and 24 h pi to quantify apoptosis and *IFNB* gene induction. The analysis of caspase 3/7 activation showed a significant reduction of apoptosis in DF-1 WT cells transfected with *MDA5* siRNA in comparison to samples that had been transfected with control siRNA, as we had previously observed (Fig 2B). However, no significant differences were identified in samples from DF-1 TLR3 KO cells transfected with *MDA5* siRNA or control siRNA at either 16 or 24 h pi (Fig 6E), as it could be expected since caspase 3/7 activity was already almost undetectable in non-transfected DF-1 TLR3 KO cells. Regarding *IFNB* gene transcription quantified by RT-qPCR, the results showed again that TLR3 deletion resulted in a significant decrease in *IFNB* mRNA levels in comparison to DF-1 WT cells (approximately 1.5 and 2 log units at 16 h pi and 24 h pi, respectively) (Fig 6F). Furthermore, the data showed a lower, but significant, decline in *IFNB* induction in both DF-1 WT and DF-1 TLR3 KO cells transfected with *MDA5* siRNA, compared to the same cells transfected with control siRNA at 16 and 24 h pi (≤1 log unit) (Fig 6F). Taken together, these results indicate a synergistic effect between MDA5 and TLR3-mediated innate immune response activation pathways in response to IBDV infection. While TLR3 appears to be the dominant contributor, MDA5 also plays a complementary role in modulating both antiviral gene expression and apoptosis.

### Involvement of TLR3 in IFN-β and NF-κB promoter activation by dsRNA

To further characterize the DF-1 TLR3 KO cells, we performed luciferase reporter assays to analyze the activation of the IFN-β and NF-κB promoters upon treatment with dsRNA in DF-1 TLR3 KO cells in the absence or presence of exogenously expressed *TLR3* gene. For this, DF-1 WT and DF-1 TLR3 KO cells were transfected with either the IFN-β or NF-κB reporter plasmids, together with a single dose of the empty pcDNA3 expression plasmid as control, or with different doses of the pc-chTLR3-His plasmid expressing TLR3 in the case of DF-1 TLR3 KO cells. At 8 h pt cells were transfected with synthetic dsRNA (poly I:C) and were harvested 16 h later. As shown in Figs. 7A-7B, when DF-1 TLR3 KO cells were transfected with the empty expression plasmid the activation of the IFN-β and NF-κB promoters was significantly lower than that found in WT DF-1 cells. However, the impairment of IFN-β and NF-κB promoter activation caused by TLR3 knockout could be rescued by TLR3 overexpression in a dose-dependent manner, reaching with the highest doses of plasmid activity levels comparable to those obtained in WT DF-1 cells. These results demonstrate that the downregulation of the innate immune response in DF-1 TLR3 KO cells is exclusively due to the absence of TLR3 protein.

**Fig 7.**
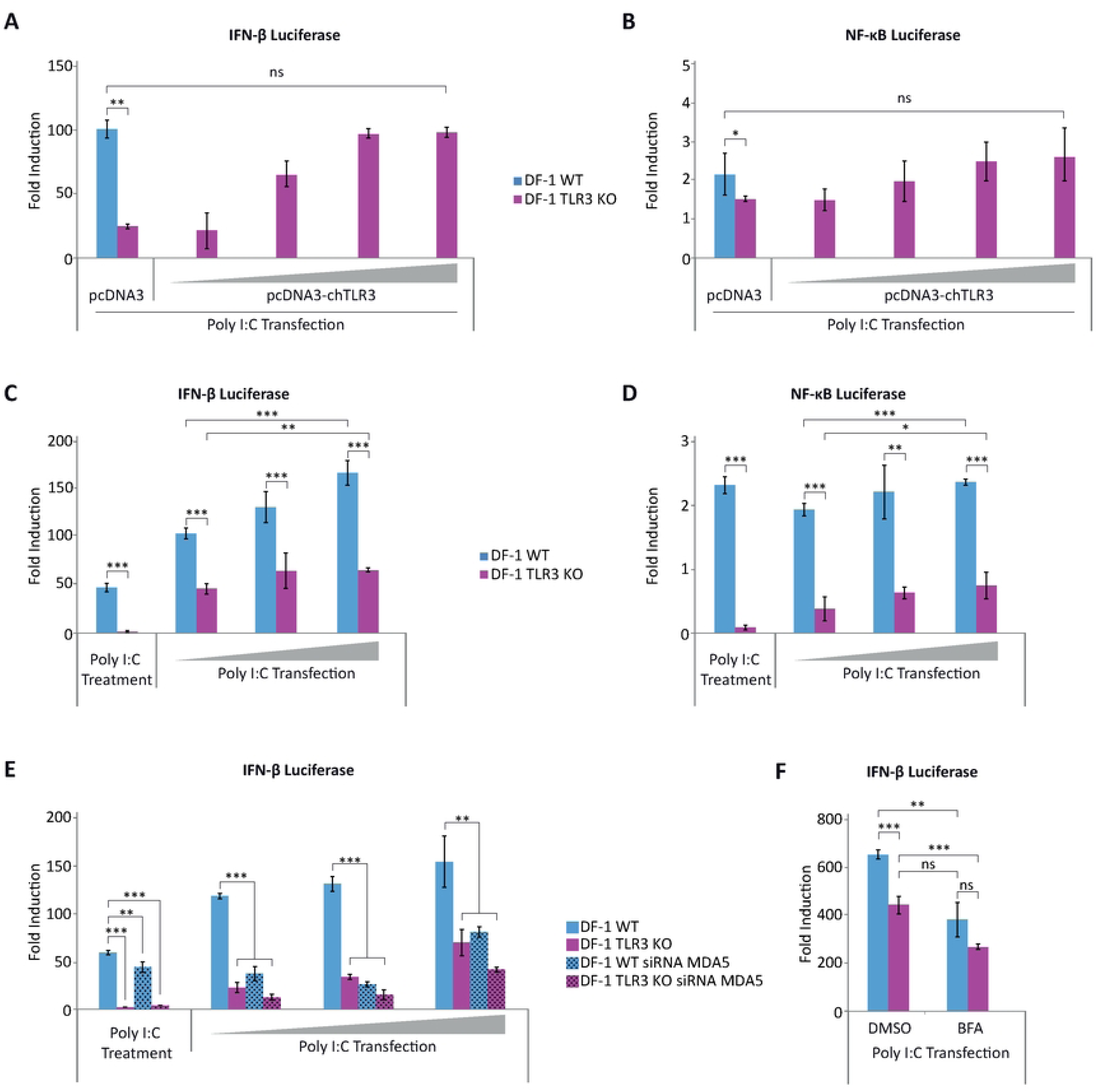
TLR3 is involved in the activation of IFN-β and NF-κB promoters by dsRNA in DF-1 cells. **(A-B)** IFN-β and NF-κB promoter activities are downregulated in DF-1 TLR3 KO cells but can be rescued by exogenous expression of TLR3. DF-1 and DF-1 TLR3 KO cells were co-transfected with pLucter (100 ng) and pR-null (30 ng) plasmids **(A)**, or with pSI-chNFκB-Luc (50 ng) plasmid **(B)** together with different amounts (100, 200, 400 or 800 ng) of the plasmid expressing chTLR3-His. At 8 h pt the cells were transfected with 250 ng of Poly I:C, and harvested at 24 h after plasmid transfection. A control consisting in cells co-transfected with pLucter, pR-null and the empty pCDNA3, or with the pSI-chNFκB-Luc, and the empty pCDNA3 plasmid (800 ng) was included in each assay. The samples were analyzed by the dual luciferase assay kit. Each determination was carried out in duplicate, and the firefly luciferase expression level of each sample was normalized by *Renilla* values. Results are expressed as the fold induction over the level found in the control DF-1 or DF-1 TLR3 KO cells not transfected with poly I:C. **(C-D)** DF-1 and DF-1 TLR3 KO cells were transfected with the plasmids pLucter (100 ng) and pR-null (30 ng) **(C)** or with the plasmid pSI-chNFκB-Luc (50 ng) **(D).** At 8 h pt, cells were either treated with Poly I:C (20 µg) added directly to the culture medium or transfected with increasing amounts (100, 200 and 300 ng) of Poly I:C. The samples were harvested at 24 h after plasmid transfection and used for luciferase activity quantification. **(E)** DF-1 and DF-1 TLR3 KO cells were transfected with a siRNA specifically silencing the expression of MDA5 protein, or with a control siRNA. At 24 h pt, the cultures were transfected with the plasmids pLucter (100 ng) and pR-null (30 ng) and 8 h later cells were either treated with poly I:C (20μg) or transfected with increasing amounts (100, 200 and 300 ng) of Poly I:C. The samples were harvested at 24 h after plasmid transfection and used for luciferase assay. **(F)** DF-1 and DF-1 TLR3 KO cells were transfected with the plasmids pLucter (100 ng) and pR-null (30 ng). At 8 h pt, cells were transfected with 250 ng of Poly I:C and subsequently treated with BFA (50μM) or the vehicle DMSO 1 h later. The samples were harvested at 24 h after plasmid transfection and analyzed using the dual luciferase assay kit. In panels C-F values are presented as the fold induction over the level found in the control DF-1 or DF-1 TLR3 KO cells not treated with Poly I:C. Bars indicate means ± standard deviations based on data of duplicate samples from three independent experiments. *, ** and *** indicate *p* values of <0.05, <0.01 and <0.001, respectively, as determined by unpaired Student’s test. ns, not significant.

In a similar set of experiments, cells were then either treated with the synthetic dsRNA poly I:C (20 μg) added directly to the culture medium or transfected with different amounts of poly I:C (100, 200 or 300 ng) at 8 h pt with the IFN-β and NF-κB reporter plasmids. 16 h later cells were harvested, and luciferase activity quantified. As shown in Fig 7, while addition of poly I:C readily activated IFN-β (Fig 7C) and NF-κB (Fig 7D) promoters in WT DF-1 cells, DF-1 TLR3 KO cells did not respond to the same treatment. Furthermore, as shown before, transfection of poly I:C into WT DF-1 cells produced a significant dose-dependent increase in IFN-β promoter activity. Again, this treatment induced some activation of the IFN-β in DF-1 TLR3 KO cells, but the highest activity attained in these cells was significantly lower (40-50%) than that in WT DF-1 cells (Fig 7C). Similarly, lower levels of NF-κB promoter activation were detected in DF-1 TLR3 KO transfected with poly I:C in comparison with WT DF-1 cells (Fig 7D).

To assess the contribution of the cytoplasmic sensor MDA5 to the activation of the IFN-β promoter by poly I:C, a similar experiment was conducted with sets of cells transfected with the *MDA5-*specific or control siRNA, and subsequently treated or transfected with poly I:C. The silencing of *MDA5* in WT DF-1 cells resulted in a slight reduction in IFN-β promoter activity in cells treated with poly I:C. As in the previous experiment, only residual activity could be detected in DF-1 TLR3 KO cells, regardless of the treatment with the *MDA5*-specific siRNA or the control siRNA (Fig 7E). However, in *MDA5*-silenced WT DF-1 cells transfected with each of the three doses of poly I:C there was a notable decline in IFN-β promoter activity in comparison with cells transfected with the control siRNA. This reduction was comparable in magnitude to that observed in DF-1 TLR3 KO cells, which was even more pronounced when these cells were transfected with the *MDA5*-specific siRNA. Overall, these results demonstrate that endosomal stimulation by poly I:C induces mainly a TLR3-dependent activation of IFN-β and NF-κB promoters. Conversely, cytoplasmic delivery of poly I:C by transfection activated both MDA5- and TLR3-signaling pathways. Activation of the TLR3-dependent pathway by transfected poly I:C suggests that the dsRNA was captured by endosomes in the cytoplasm.

To further confirm the role of TLR3 in the detection of transfected poly I:C, we used bafilomycin A1 (BFA) to prevent endosomal acidification, which is required for TLR activation (Hacker et al., 1998; Yi et al., 1998). Therefore, DF-1 WT and DF-1 TLR3 KO previously transfected with the IFN-β reporter plasmid were transfected with poly I:C (250 ng) and subsequently treated with BFA (50 μM) or the vehicle dimethyl sulfoxide (DMSO) 1 h later. Following a 16 h incubation period, the cells were harvested, and the luciferase activity quantified. Fig 7F shows that treatment with BFA of poly I:C-transfected WT DF-1 cells resulted in a notable reduction (42%) in IFN-β promoter activity, approaching the result obtained in DF-1 TLR3 KO cells in the absence of BFA. Significantly, a reduction in IFN-β promoter activity was also observed upon treatment of DF-1 TLR3 KO cells with BFA, indicating that in addition to TLR3, other TLRs could also be activated in IBDV-infected cells.

### Endosomal acidification is required for TLR3-mediated apoptosis induction in IBDV-infected DF-1 cells

Given that endosomal acidification is a prerequisite for TLR3-dependent signaling, we sought to ascertain the impact of the BFA treatment on the induction of apoptosis in response to IBDV infection. Therefore, DF-1 WT and DF-1 TLR3 KO cells infected with IBDV were treated with BFA or the vehicle DMSO 2 h later and caspase 7/3 activity was analyzed at 24 h pi. As observed in previous experiments, a significant reduction in luciferase levels was evident in DF-1 TLR3 KO cells in comparison with WT DF-1 cells. It is noteworthy that treatment of WT DF-1 cells with BFA resulted in a marked reduction in caspase 3/7 activity, with luciferase values comparable to those observed in DF-1 TLR3 KO cells (Fig 8A). These findings further substantiate the pivotal role of TLR3 in apoptosis induction by IBDV infection. To confirm that BFA does not affect IBDV replication under the conditions used, we conducted an RT-qPCR analysis to quantify the accumulation of viral RNA in samples from cells treated as described above. As illustrated in Fig 8B, the RNA accumulation values were markedly higher in DF-1 TLR3 KO cells compared to WT DF-1 cells, regardless of BFA treatment. Neither BFA treatment affected the accumulation of IBDV RNA in WT DF-1 cells. We also analyzed the production of intra- and extracellular infectious virus by virus titration from both, cell supernatants and extracts from infected cell cultures. As shown previously, the accumulation of intracellular virus was higher in DF-1 TLR3 KO cells, and BFA treatment did not alter the rate of virus accumulation in either cell line (Fig 8C). On the other hand, the titer of extracellular virus was again significantly higher in WT DF-1 cells compared to DF-1 TLR3 KO cells, and BFA treatment produce a slight, not significant, reduction in virus titer in WT DF-1 cells. A more pronounced effect was observed in BFA-treated DF-1 TLR3 KO cells (Fig 8D). A possible explanation for the reduction in extracellular virus titers could be the presence of BFA in the supernatants of BFA-treated cells, which could interfere with endocytic virus entry into the cells of the cultured monolayers used for titration. Indeed, pretreatment of DF-1 cells with BFA for 30 min before infection causes a five to ten-fold reduction in both extracellular and intracellular virus yields as determined at 24 h pi (data nor shown).

**Fig 8.**
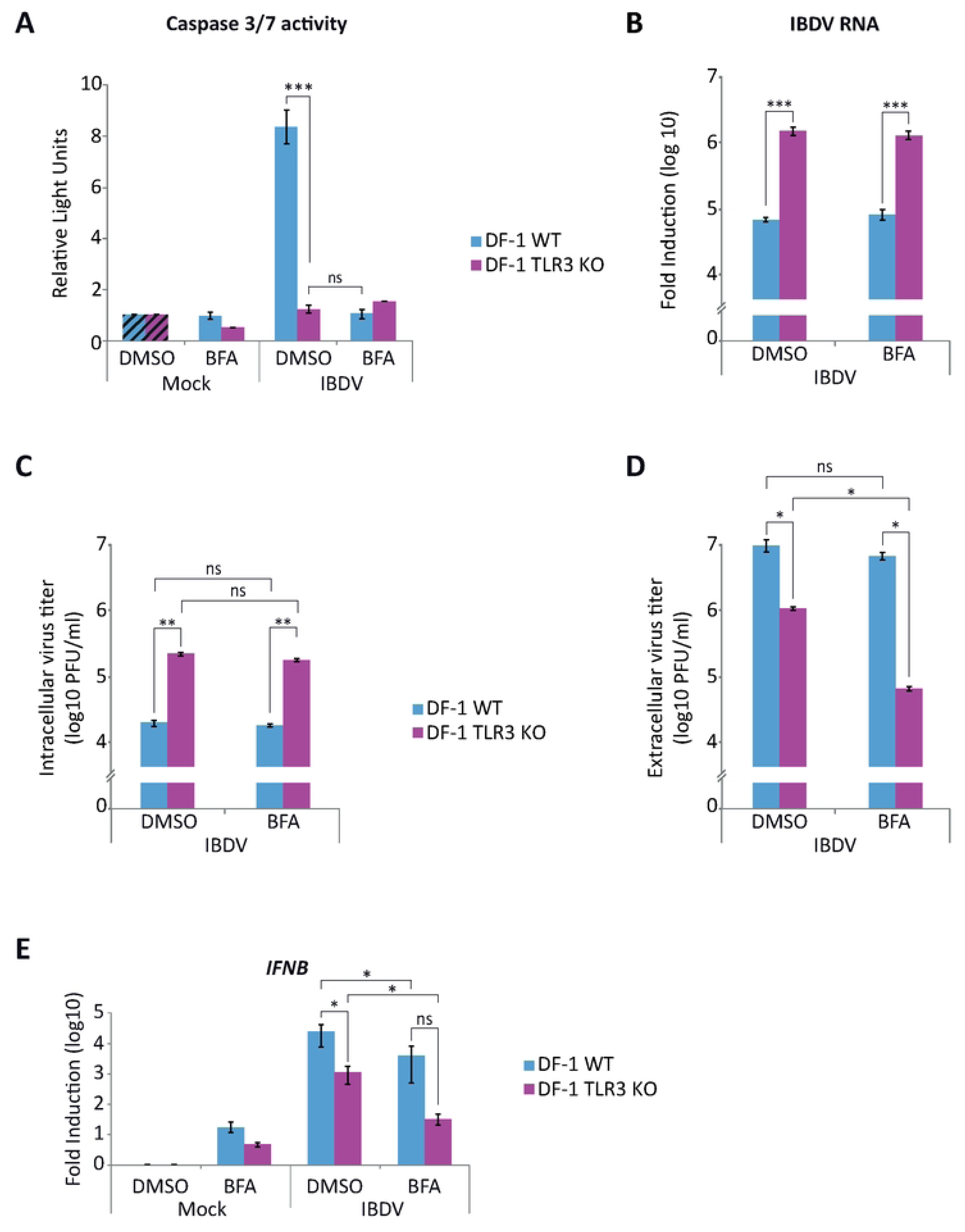
Inhibition of endosomal acidification by treatment of IBDV-infected DF-1 cells with BFA abrogates apoptotic cell death. DF-1 and DF-1 TLR3 KO cells mock-infected or infected with IBDV (MOI of 2 PFU/ml) were treated or not with BFA (50 µM) after 2 h pi. **(A)** Apoptosis was measured by using the Caspase-Glo 3/7 assay kit, and each determination was carried out in duplicate. Caspase values from infected cell samples were normalized to those from mock-infected cells (Mock) (Striped bars). **(B)** The accumulation of IBDV RNA was determined by SYBR green-based RT-qPCR. Analysis of extracellular **(C)** and intracellular **(D)** virus titers. **(E)** RT-qPCR study of the expression of *IFNB* gene. Cellular gene expression values were normalized to those of the *GAPDH* mRNA content and are presented on a log_10_ scale as the fold induction over the level found in mock-infected DF-1 or DF-1 TLR3 KO cells not treated with BFA. Bars indicate means ± standard deviations based on data of duplicate samples from three independent experiments. *, ** and *** indicate *p* values of <0.05, <0.01 and <0.001, respectively, as determined by unpaired Student’s test. ns, not significant.

*IFNB* gene expression was also analyzed in samples from BFA treated WT DF-1 and DF-1 TLR3 KO cells. In BFA-untreated samples we observed a diminished expression in DF-1 TLR3 KO cells as compared to DF-1 WT cells, as previously shown. An additional reduction was observed in both DF-1 WT and DF-1 TLR3 KO cells treated with BFA. The decrease of *IFNB* gene expression in BFA-treated DF-1 TLR3 KO could again indicate that other TLR can also be activated in IBDV-infected cells. Although the effect of BFA treatment on *IFNB* gene expression in DF-1 WT cells was clear and statistically significant, it was not as drastic as the effect on apoptosis (compare Figs. 8A and 8E). This result suggests that TLR3 deficiency has a direct effect on apoptosis, in addition to its indirect effect through the reduction of *IFNB* gene expression.

### Role of RIPK1 in IBDV-mediated apoptosis triggering

About two decades ago, several laboratories described that TLR3 can directly trigger caspase-8-dependent apoptosis in human cancer cells upon activation by dsRNA, a process that requires the formation of a complex involving caspase 8, TLR3 and TRIF [39,40]. Later, it was demonstrated that recruitment and activation of caspase 8 to TLR3 requires the participation of RIPK1 [41]. In view of our previous results suggesting a direct role of TLR3 in apoptosis triggering in IBDV-infected cells, additionally to its contribution through IFN-β production, we sought to investigate the potential involvement of RIPK1 in this apoptotic process. After multiple unsuccessful attempts to significantly knock down RIPK1 expression using siRNA, we employed CRISPR/Cas9 gene editing to generate a large deletion (∼15.7 kb) in the *RIPK1* gene of DF-1 cells, spanning from exon 2 to exon 12, the last exon of the gene (Fig 9A). Two clones, designated 9 and 12, displaying minor variations in deletion length, were selected from the pool of genetically characterized clones for further analysis. First, we evaluate the fate of RIPK1 deficient DF-1 cells following IBDV infection. For this, DF-1 WT and DF-1 RIPK1 KO cells from both clones were infected and samples were taken at 16 and 24 h pi for analysis of caspase 3/7 activity. Strikingly, a marked reduction in apoptosis levels was observed in infected cells from both RIPK1 KO cell lines compared to DF-1 WT cells at both pi time points (Fig 9B). This effect was particularly pronounced at 24 h pi, when apoptosis levels in DF-1 WT cells were notably higher. Remarkably, as it occurred with DF-1 TLR3 KO cells, the extent of apoptosis in both cell clones following IBDV infection was comparable to that measured in uninfected control cells. With respect to DF-1 WT cells, the expression of *IFNB* gene in the absence of RIPK1 was also reduced at both times pi for each of the two clones analyzed (Fig 9C), although the differences were only statistically significant at 24 h pi. Moreover, the reduction of *IFNB* gene expression was below 1-log unit, which is lower than that observed in the DF-1 TLR3 KO cells (compare Figs. 6A and 9B). Accordingly, only minor differences in the accumulation of viral RNA between WT DF-1 and DF-1 RIPK1 KO cells were observed at the two times pi analyzed (Fig 9D). Overall, these findings indicate that RIPK1 is a key mediator of apoptosis induction in DF-1 cells following IBDV infection, while also contributing partially to IFN-β signaling, as evidenced by the modest reduction in *IFNB* gene expression observed in RIPK1-deficient cells.

**Fig 9.**
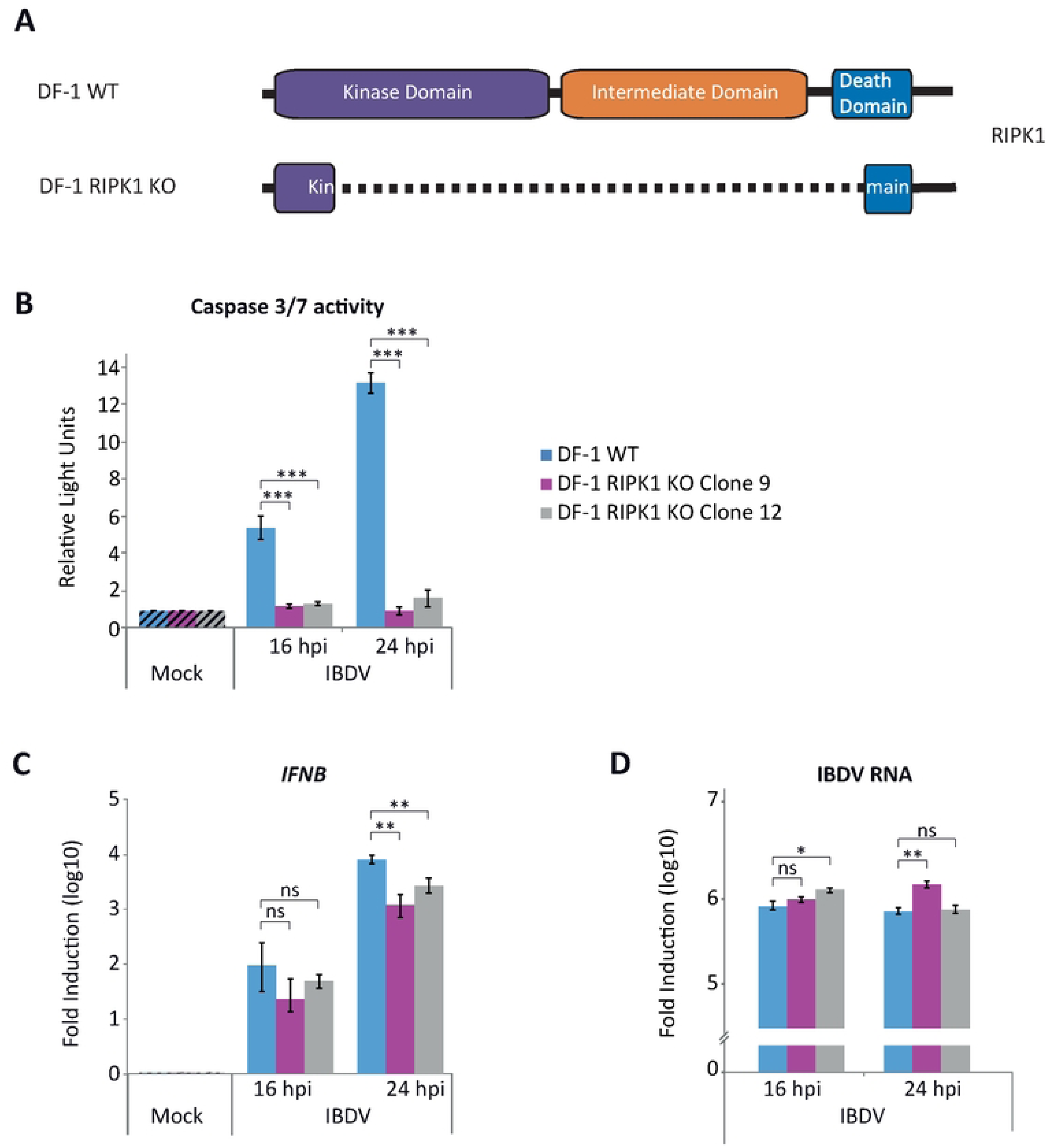
Ablation of RIPK1 expression abrogates IBDV-mediated apoptosis triggering. **(A)** Schematic representation of the structure of the *RIPK1* gene and the deleted versions of the gene present in two selected DF-1 RIPK1 KO cell clones. **(B)** DF-1 and DF-1 RIPK1 KO cells from clones 9 and 12 were mock-infected or infected with IBDV (MOI of 2 PFU/cell) and cells were harvested at 16 and 24 h pi. Apoptosis was measured by using the Caspase-Glo 3/7 assay kit, and each determination was carried out in duplicate. Caspase values from infected cell samples were normalized to those from mock-infected cells (Mock) (Striped bars). Bars indicate means ± standard deviations based on data of duplicate samples from three independent experiments. *** indicate *p* value of <0.001, as determined by unpaired Student’s test. **(C-D)** DF-1 and DF-1 RIPK1 KO cells mock-infected or infected with IBDV (MOI of 2 PFU/cell) were harvested at 24 h pi and the expression levels of the *IFNB* gene **(C)** and the accumulation of viral RNA **(D)** were analyzed by SYBR green-based RT-qPCR. Recorded values for *IFNB* gene were normalized to the *GAPDH* mRNA content and are presented on a log_10_ scale as the fold induction over the level found in mock-infected DF-1 or DF-1 RIPK1 KO cells. Bars indicate means ± standard deviations based on data of duplicate samples from three independent experiments. * and ** indicate *p* values of <0.05 and <0.01, respectively, as determined by unpaired Student’s test. ns, not significant.

### TLR3 deletion does not affect replication of avian reovirus (ARV), vesicular stomatitis virus (VSV), Semliki Forest virus (SFV) and Newcastle disease virus (NDV)

The enhancement of IBDV replication and its intracellular accumulation in cells that do not express TLR3 is consistent with data supporting the involvement of TLR3 in the regulation of the innate immune response and in the activation of apoptosis induced by IBDV infection. We therefore decided to investigate whether this effect is specific to IBDV infection or if the absence of TLR3 also affects the replication of other RNA viruses. To this end, DF-1 WT and DF-1 TLR3 KO cells were infected with ARV (another dsRNA virus), VSV-GFP (a negative polarity ssRNA virus expressing GFP), and SFV (a positive polarity ssRNA virus) at an MOI of 2 PFU/cell, and the extra- and intracellular viral titers quantified at 16 and 24 h pi. Both intra- and extracellular titers of ARV (Fig 10A), VSV-GFP (Fig 10B) and SFV (data not shown) did not vary depending on the infected cell line, DF-1 WT or DF-1 TLR3 KO. Additionally, we assessed NDV-GFP replication in both DF-1 WT and DF-1 TLR3 KO cell lines by quantifying GFP fluorescence. Cells were infected at MOIs of 0.1 and 1 PFU/cell, and samples were collected at 24 h pi. No significant differences in GFP expression were observed between the two cell lines at either MOI tested (Fig 10C).

**Fig 10.**
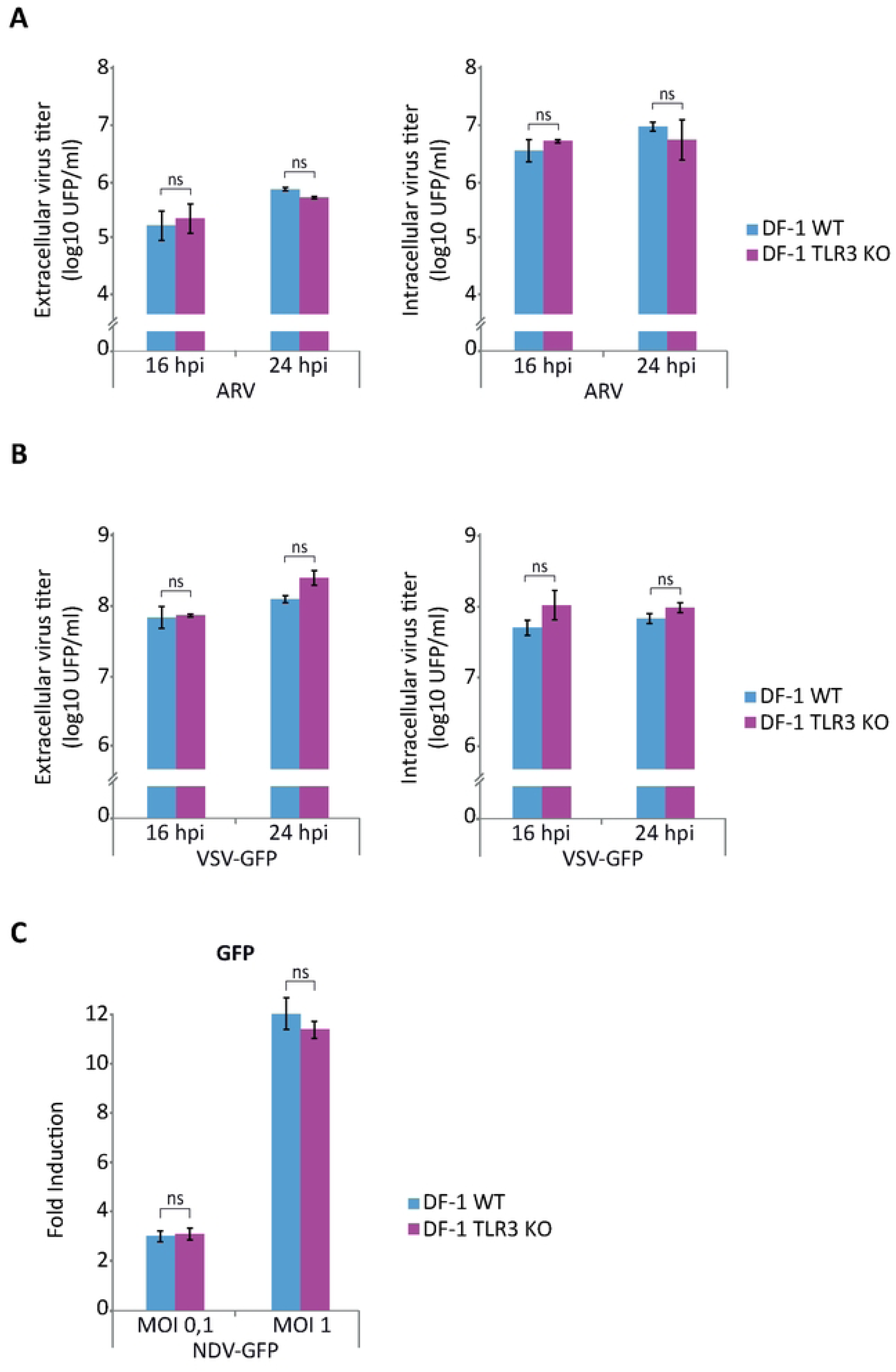
TLR3 deletion does not influence ARV, VSV-GFP and NDV-GFP replication. **(A-B)** DF-1 and DF-1 TLR3 KO cells were mock-infected or infected with ARV **(A)** and VSV-GFP **(B)** (MOI of 2 PFU/cell) and cells were harvested at 16 and 24 h pi for analysis of extracellular and intracellular virus titers. **(C)** DF-1 and DF-1 TLR3 KO cells were mock-infected or infected with NDV-GFP at MOIs of 0.1 and 1 PFU/cell and cells were harvested at 24 h pi for analysis of GFP expression. Recorded values for GFP fluorescence were normalized to those found in mock-infected DF-1 or DF-1 RIPK1 KO cells. Bars indicate means ± standard derivations based on duplicate samples from three independent experiments. ns, not significant.

## DISCUSSION

IBDV represents a major challenge to the host immune defense in juvenile chickens, as it targets and eliminates developing B lymphocytes within the bursa of Fabricius, their primary lymphoid organ responsible for B cell maturation and the establishment of a diverse antibody repertoire in birds. The massive depletion of B lymphocytes due to apoptotic cell death causes an irreversible atrophy of this organ, resulting in profound and long-lasting immunosuppression [42]. Different studies have provided evidence for the implication of IBDV proteins in the apoptotic process. Specifically, VP2 was first identified as an apoptotic inducer ([43,44], but later studies have revealed that VP5 also promotes apoptosis [45,46]. Our previous results have shown that exposure to type I IFN in IBDV-infected cells induces massive cell death through apoptosis, suggesting that IFN may play a critical role in the pathogenesis associated with this virus.

In this study, we demonstrate that treatment of chicken DF-1 cells with chIFN-α shortly after IBDV infection not only accelerates the expression of the *IFNB* gene, as well as those of all tested ISGs, but also significantly anticipates the onset of cytopathic effects associated with apoptotic cell death. This was evidenced by the earlier activation of caspases 3/7 (16 h pi) compared to untreated infected cultures, in which significant caspase activity was only detected at 24 h pi. Conversely, pharmacological inhibition of type I IFN signaling through blockade of the JAK/STAT pathway by treatment with Rx— which markedly suppresses the innate immune response, as indicated by reduced expression of *IFNB*, *MDA5*, *TLR3*, and *Mx* genes—also has a profound impact on the outcome of infection, preventing apoptotic cell death. This was reflected by a dramatic reduction in caspase 3/7 activity at 24 h pi in IBDV-infected cells treated with the inhibitor, indicating that endogenous type I IFN production contributes to apoptosis at later stages of infection. Additionally, inhibition of IBDV replication also causes a drastic drop in caspases 3/7 activity. These findings are consistent with the notion that IBDV dsRNA is the viral factor responsible of apoptosis triggering, as previously indicated by work performed in HeLa cells [16]. In this regard, previous studies showed that recognition of IBDV genomic dsRNA by MDA5 activates the MAVS-dependent signaling cascade to induce type I IFN expression [20–23]. Consistently, our results show that individual or combined silencing of *MDA5* and *MAVS* genes in DF-1 cells prior to infection leads to attenuated *IFNB* gene expression, and results in a significant, albeit partial, reduction of apoptosis induction and enhanced IBDV replication. Altogether, these findings highlight the pro-apoptotic role of type I IFN during IBDV infection and support the notion that, beyond establishing an antiviral state, IFNs actively promote programmed cell death in infected cells as an additional mechanism to limit viral spread, as previously reported in other viral infection models [47–48].

Several studies have confirmed the sensitivity of IBDV to the antiviral action of IFNs, with both type I and II IFNs shown to reduce viral replication *in vitro* and *in vivo* [16, 49,50]. However, our findings are consistent with previous studies that link type I IFN responses with the severity of IBDV-induced pathology. Pathogenic and very virulent IBDV strains have been shown to induce stronger or dysregulated IFN responses, which correlate with increased bursal damage and mortality [51–53]. Moreover, it was reported that more susceptible chicken breeds mount earlier and more robust IFN responses following infection [53]. Together, these studies highlight that while IFNs are essential for antiviral defense, excessive or early activation may enhance immunopathology and contribute to disease progression. The dual, context-dependent role of type I IFNs has been well documented in various viral infection models, such as human immunodeficiency virus (HIV), lymphocytic choriomeningitis virus (LCMV), and severe acute respiratory syndrome coronavirus (SARS-CoV), where systemic IFN responses contribute to either immune suppression or enhanced pathology [54–56].

In view of the significant abrogation of apoptosis in IBDV-infected cell cultures following inhibition of the JAK/STAT pathway by Rx treatment and considering the feasibility of its *in vivo* administration [57], this compound may constitute a valuable experimental tool for elucidating the functional significance of IFN signaling in the pathogenesis of IBDV under physiological conditions.

### PKR contributes to apoptosis in IBDV-infected cells, although it is not the primary driver of this process

As mentioned in the introduction, PKR is an ISG that plays a central role in the host antiviral defense and the regulation of apoptosis. PKR is also a dsRNA cytoplasmic sensor, which becomes activated upon binding to viral or synthetic dsRNA, initiating downstream signaling cascades that inhibit protein translation and promote cell death [18,58,59]. Our previous work revealed that PKR plays a pivotal role in apoptosis induction in human HeLa cells infected with IBDV and treated with IFN-α, a process that is dependent on the presence of IBDV dsRNA [16]. Upregulation of PKR, as well as of different ISGs, has been shown to occur not only in cells infected *in vitro* [29,60], but also in the bursa of Fabricius and the spleen following IBDV infection [28]. Then, to investigate the potential role of PKR in IBDV-induced apoptosis in chicken cells we generated DF-1 PKR KO cells. Our results show that ablation of PKR expression significantly prevents the cell damage inflicted by IBDV infection in chicken cells. In parallel, a reduction in *IFNB* gene expression was also observed in these cells, suggesting that as it has been described for its mammalian ortholog, chicken PKR is required for maximal induction of type I IFN [18,61]. Nonetheless, the remaining apoptotic activity observed in PKR-deficient cells indicates that alternative signaling pathways or mediators are likely to participate in the execution of IBDV-induced apoptosis in chicken cells.

### TLR3 plays a crucial role in IFN expression and apoptosis in IBDV-infected DF-1 cells

Results discussed above evidence that reducing the expression of the cytoplasmic sensor MDA5 or its adaptor MAVS, as well as deleting PKR, consistently lead to a decrease of IFN-β expression in IBDV-infected DF-1 cells and diminished levels of apoptosis, reinforcing the contribution of these cytosolic pathways to the antiviral and pro-apoptotic responses. However, in none of these cases IFN-β expression was fully abolished, nor was apoptosis entirely prevented, suggesting that additional pathways contribute to these processes.

Several studies have proposed that the endosomal receptor TLR3 might be involved in the recognition of IBDV dsRNA [27,62]. Experimental evidence showed that TLR3 expression is upregulated during IBDV infection both *in vitro* and *in vivo*, supporting its potential role in the innate immune response to the virus [28,30,63]. Furthermore, a correlation has been observed between TLR3 levels in the bursa of Fabricius and both IFN-β expression and the virulence of the IBDV strain used for the experimental infection, suggesting the hypothesis that more virulent strains might elicit stronger TLR3-mediated responses [26]. Notably, Guo et al. [27] demonstrated that TLR3 is capable of recognizing the synthetic dsRNA analog poly I:C in vaccinated chickens and proposed that this recognition pathway might also be activated by IBDV-derived dsRNA, thus linking TLR3 activation with downstream interferon production during infection (Guo et al., 2012). However, despite all this supporting data highlighting a potentially important role for TLR3 during IBDV infection, direct demonstration of TLR3’s involvement in IFN-β production in IBDV-infected cells remained to be experimentally validated.

To explore the role of TLR3 during IBDV infection we generated DF-1 TLR3 KO cell lines using the CRISPR/Cas9 technology. Strikingly, microscopic examination of IBDV-infected DF-1 TLR3 KO cells at late pi stages revealed a complete absence of the cytopathic effects typically observed in infected WT DF-1 cells. Caspase 3/7 activity assays confirmed that apoptotic activity in TLR3-deficient cells remains at basal levels, indistinguishable from those of mock-infected controls. Notably, this resistance to apoptosis is accompanied by a marked reduction in the expression of *IFNB*, *MDA5, Mx* and *OAS* genes, pointing TLR3 as the central mediator of both antiviral signaling and apoptosis in IBDV-infected chicken cells. Nonetheless, the observation that knocking down MDA5 expression in DF-1 TLR3 KO cells further reduces the *IFNB* gene expression induced by IBDV infection, indicates a synergistic effect of both PRRs orchestrating the innate immune response.

Transactivation reporter assays conducted in DF-1 TLR3 KO cells further underscored the pivotal role of chTLR3 in the activation of IFN-β and NF-κB promoters in response to dsRNA stimulation. The impaired activation of both reporter elements in TLR3-deficient cells upon transfection with poly I:C is rescued in a dose-dependent manner by exogenous expression of chTLR3 from a plasmid vector. This experimental system enabled the evaluation of the relative contributions of TLR3 and MDA5 to the transactivation of the IFN-β and NF-κB promoters upon exposure to poly I:C, either via extracellular (endosomal) treatment or intracellular (cytoplasmic) transfection. While DF-1 WT cells responded to stimulation by either route, the addition of poly I:C to the culture medium failed to activate both promoters in TLR3 KO cells. However, *MDA5* silencing had only a minor impact in IFN-β promoter activation in DF-1 WT cells treated with poly I:C. In contrast, when poly I:C was delivered by transfection, both TLR3 deletion and MDA5 knockdown independently reduced IFN-β promoter activity, with a more pronounced decrease observed upon combined inhibition of both PRRs. These findings indicate that extracellular poly I:C predominantly activates TLR3-dependent signaling pathways correlating with its endocytic uptake, whereas cytoplasmic delivery engages both MDA5- and TLR3-mediated responses. The observation that transfected poly I:C can also trigger TLR3 signaling suggests that cytoplasmic dsRNA may be entrapped into endosomes, a mechanism previously reported in HeLa cells microinjected with poly I:C [64]. In agreement with this interpretation, inhibition of endosomal acidification by treatment of poly I:C transfected DF-1 WT cells with BFA results in a notable reduction in IFN-β promoter activity. To our surprise, we also observed a clear, although not statistically significant, reduction in IFN-β promoter activity in DF-1 TLR3 KO cells after treatment with BFA. This result could indicate that poly I:C transfection may also activate other endosomal receptors, such as TLR7, whose expression is also upregulated after IBDV infection [26,28].

Notably, inhibition of endosomal acidification during IBDV infection completely abrogated apoptosis induction in DF-1 WT cells, closely mimicking the results obtained in DF-1 TLR3 KO cells. This was accompanied by a reduction in *IFNB* gene expression, although the extent of this decrease was less pronounced than that observed for caspase 3/7 activity. A minor effect in *IFNB* gene expression was also observed in BFA-treated DF-1 TLR3 KO cells, suggesting again the potential activation of TLR7. Importantly, it was assessed that the BFA treatment (applied 3 h after virus adsorption) did not affect intracellular viral RNA accumulation or virus production, discarding the possibility that the observed effects could result from inhibition of IBDV infection. Only a slight reduction in extracellular viral titers from treated DF-1 TLR3 KO cells was noted, which could be attributable to residual BFA in the culture supernatants used for titration, potentially impairing endocytic uptake of virus during the plaque assay. Supporting this notion, pretreatment of DF-1 cells with BFA for 30 minutes prior to infection resulted in a five- to ten-fold decrease in both extracellular and intracellular virus yields at 24 h pi. The pronounced effect of either TLR3 ablation or pharmacological inhibition of TLR3 activation on apoptosis induction—contrasted with the more moderate reduction observed in *IFNB* gene expression—suggests that TLR3 may play functionally distinct roles during IBDV infection in DF-1 cells. These findings imply that TLR3-mediated cell death is not solely a consequence of its canonical function in initiating the IFN signaling cascade, but also reflects a more direct, IFN-independent contribution to the regulation of apoptosis.

### Role of TLR3 in the control of viral replication and spread

Regarding the role of TLR3 on viral replication, the results obtained in DF-1 TLR3 KO cells showed an increased accumulation of VP3 protein and viral RNA, but surprisingly a decrease in extracellular viral titers was observed. We therefore decided to analyze intracellular viral titers and observed an increase in the levels achieved in DF-1 TLR3 KO cells compared to the parental cell line. These results suggest that the absence of TLR3 favors viral replication, but also has a negative impact on viral shedding into the extracellular medium. This effect could be related to the inhibition of apoptosis in DF-1 TLR3 KO cells and/or a blockade of the viral exit pathway. The first possibility contrasts with our previous results in which we observed that reduced apoptosis was always accompanied by an increase in extracellular virus titers, as it has been shown for other RNA viruses such as Sendai virus (SeV) and Sindbis virus (SINV) [64]. On the other hand, the hypothesis that deletion of the *TLR3* gene affects viral exit is supported by experiments with a spreading deficient IBDV mutant virus. Previous results from our laboratory have shown that IBDV uses a non-lytic egression mechanism dependent on the VP5 protein during early stages of infection [7,8]. The extracellular virus titers obtained with the mutant virus lacking the VP5 protein (IBDV VP5 KO) do not vary depending on the infected cell line, DF-1 WT or DF-1 TLR3 KO, and are similar to those obtained with IBDV WT in the DF-1 TLR3 KO cell line. These results suggest that the TLR3 protein and the viral protein VP5 may act in a coordinated manner in the egression of viral particles from cells. Further research will be required to understand the underlying mechanism involved in TLR3-dependent IBDV release. On the other hand, as with the WT virus, higher intracellular virus titers were obtained with IBDV VP5 KO from DF-1 TLR3 KO cells. Again, the increase in IBDV intracellular virus titer in DF-1 TLR3 KO cells correlates with decreased activation of the IFN-β production pathway in these cells (data not shown).

When analyzing the effect of the lack of TLR3 expression on infection with other viruses with RNA genomes of different types, such as ARV (dsRNA), VSV (single-stranded RNA and negative polarity), SFV (single-stranded RNA and positive polarity) or NDV (single-stranded and negative polarity), which enter the cells via the endocytic pathway, we did not observe significant differences in the intracellular and extracellular viral titers obtained from DF-1 WT and DF-1 TLR3 KO cell lines. These results align with those reported for ARV, VSV, and influenza A virus (IAV) in quail fibroblast cells lacking TLR3. In that study, it was shown that TLR3-deficient cells failed to induce IFN-β expression following stimulation with poly(I:C). However, during infection with the aforementioned viruses, no significant differences in the expression of immune-related genes were observed between WT and TLR3 KO, and consequently each virus replicates similarly in WT and TLR3 KO cell lines [65]. In DF-1 cells, MDA5 has been identified as the primary sensor responsible for recognizing IAV and synthetic dsRNA, thereby initiating the innate immune response. In contrast, TLR3 appears to play a secondary role in the detection of both IAV and poly I:C in this cellular context [66]. On the other hand, a recent study by Lee and colleagues [67] in DF-1 cells showed that TLR3 deletion has a positive effect on NDV replication, although a double deletion of MDA5 and TLR3 has a greater effect. However, the authors were unable to establish a direct relationship between IFN expression and NDV replication [67]. In light of these findings, we infected DF-1 WT and DF-1 TLR3 KO cells with a recombinant NDV-GFP virus and found comparable GFP expression levels in both cell types, suggesting that TLR3 does not play a major role in restricting NDV replication under our experimental conditions. Although the experimental approaches differed between studies, our results are consistent with the notion that NDV replication is largely independent of TLR3-mediated signaling. Interestingly, Lee et al. [67] also reported that treatment with poly(I:C) induced a cytopathic effect that they described as “cell degeneration” in both DF-1 WT and MDA5 KO cells, but not in TLR3 KO cells. Although the nature of this effect was not further characterized to confirm whether it corresponded to apoptotic cell death, their findings are fully consistent with our results demonstrating that chicken TLR3 plays a pivotal role in mediating apoptosis in response to viral or synthetic dsRNA. To our knowledge, this study offers the most compelling evidence to date for the antiviral function of chicken TLR3 during IBDV infection, as well as its critical involvement in the induction of apoptosis. These findings highlight the dual role of TLR3 in orchestrating both innate immune responses and the regulation of virus-induced cell death.

### Role of RIPK1 downstream of TLR3 in the induction of apoptosis in IBDV-infected DF-1 cells

Our findings underscore a dual role for TLR3 in IBDV pathogenesis: as a sensor of viral dsRNA that initiates type I IFN responses, and as a trigger of programmed cell death. This functional profile is consistent with reports from mammalian systems, where TLR3 has been shown to elicit protective immune responses against dsRNA viruses such as poliovirus (PV), coxsackievirus (CV), and encephalomyocarditis virus (EMCV), as well as certain DNA viruses like herpes simplex virus 1 (HSV-1) [68]. However, beyond its antiviral role, several studies have shown that TLR3 activation by dsRNA also initiates apoptotic pathways in various human tumor cell lines, a process that is dependent on the recruitment and activation of caspase 8 via a signaling complex involving TRIF and RIPK1 [69]. To determine whether these molecular events are conserved in avian systems—specifically in the context of TLR3-mediated apoptosis during IBDV infection—here we generated and genetically characterized DF-1 RIPK1 KO cells. Significantly, ablation of RIPK1 expression in DF-1 cells has a dramatic effect on IBDV-induced apoptosis as determined by the reduced caspase 3/7 activity levels at late times pi with IBDV, that were equal or slightly higher than those detected in mock-infected cell cultures. A minor reduction in the level of induction of the *IFNB* gene was observed in cells lacking RIPK1, although the differences with WT cells at 24 h pi were statistically significant. Accordingly, only a slight enhancement in viral RNA accumulation was attained in DF-1 RIPK KO cells. Based on these results, we conclude that RIPK1 plays a central role in mediating apoptosis in DF-1 cells following IBDV infection, while having only a minor influence on type I IFN expression and viral replication. The near-complete abrogation of caspase activity in RIPK1-deficient cells highlights its critical contribution to TLR3-driven pro-apoptotic signaling. These results provide the first experimental evidence of the essential role of RIPK1 in virus-induced cell death in avian systems and highlight the importance of TLR3-RIPK1 signaling in shaping the pathogenesis of IBDV through modulation of host cell fate.

Our previous work in HeLa cells demonstrated that IBDV genomic dsRNA is a major determinant of apoptosis induction, with PKR playing a pivotal role in mediating this process. In contrast, the present study in chicken DF-1 cells reveals that while PKR also contributes to apoptosis upon IBDV infection, TLR3 emerges as the principal effector. Notably, ablation of TLR3 expression alone is sufficient to completely prevent IBDV-induced cell death. Recently, Zuo and collaborators have described that intracytoplasmic injection or transfection of synthetic dsRNA in HeLa cells induce massive apoptosis, which is further enhanced by type I IFN priming [64]. Contrasting with our results in DF-1 cells, they reported that PKR and TLR3 are both essential for full induction of dsRNA-mediated apoptosis in these cells. Collectively, these results suggest that while dsRNA-triggered apoptotic pathways are conserved across species, the molecular dependencies differ between human and avian systems—likely reflecting evolutionary divergence in the architecture of innate immune signaling networks.

Furthermore, our findings provide new insights into the mechanism of induction of the innate immune response during IBDV infection and its close relationship with the activation of the apoptotic death process. The results support a model in which, following infection, the viral genome is primarily recognized by the TLR3 receptor after capsid disassembly at the endosome promoted by low calcium concentration and acidic pH. The rupture of the endosomal membrane by capsid-associated amphipathic peptides, including pep46, enable the release of viral genome into the cytoplasm, where viral transcription and replication processes take place. The newly synthesized dsRNA is recognized by the MDA5 receptor and PKR protein, leading to the induction of IFN-β gene expression and apoptosis. Additionally, our results and those of Zou and collaborators [64] indicate that cytoplasmic dsRNA can be entrapped into endosomes and be sensed by TLR3, which in turn would activate two parallel signaling pathways, the one leading to IFN-β production, and the RIPK1-dependent pathway leading to apoptotic cell death. Notably, IFN-β secreted at late times pi further contributes to cell killing. This work provided the first direct link between the TLR3 receptor and the activation of the innate response to IBDV, emerging as the main receptor involved in IBDV recognition. Our results also show that TLR3 plays a crucial role on the release of the IBDV progeny which should be the subject of future studies.

TLR3 may exert contradictory roles in the context of different viral and immune-mediated diseases, such as cancer, autoimmune disorders, and allergies, by orchestrating both protective immune functions and pathological effects [70]. Specifically, in the context of infections TLR3 can enhance anti-viral defenses and promote pathogen clearance, contributing to host protection. However, evidence for a detrimental contribution of TLR3 in viral pathogenesis emerged from studies in mouse models and in certain viral infections in humans, which show that TLR3 can exacerbate tissue damage and facilitate viral replication [71]. Our results seem to be in line with the adverse effects of TLR3 on the course of IBDV infection being responsible for inducing massive cell death of infected cells. Nonetheless, further research in experimental animals would be required to determine its potential involvement in IBDV pathogenesis.

## Materials and methods

### Cells, viruses and infections

DF-1 (Chicken embryonic fibroblasts; ATCC CRL-12203) and QM7 (quail muscle myoblasts, ATCC CRL-1962) lines were grown in Dulbeccós modified minimal essential medium (DMEM) supplemented with penicillin (100 U/ml), streptomycin (100 U/ml), gentamicin (50 µg/ml), fungizone (12 µg/ml), nonessential amino acids, and 10% fetal calf serum (FCS; Sigma-Aldrich).

The WT and VP5-KO IBDV viruses used in this report are derivatives of the IBDV Soroa strain, a cell-adapted, serotype I virus, generated by reverse genetics in our laboratory (Méndez et al., 2015). IBDV infections were performed on preconfluent (70-80%) cell monolayers diluted in DMEM at an MOI of 2 PFU/cell, unless otherwise stated. After 1 h of adsorption at 37°C, the medium was removed and replaced with fresh DMEM supplemented with 2% FCS. Infected cells were incubated at 37°C until the specified times pi.

Infections with VSV-GFP (Ostertag et al., 2007), ARV strain S1133 (Kindly provided by Dr. José Manuel Martínez Costas, CIQUS, University of Santiago de Compostela, Spain), SFV (Kindly provided by Dr. Christian Smerdou, CIMA, Universidad de Navarra, Pamplona) were performed as described above on preconfluent (70-80%) DF-1 cell monolayers at an MOI of 2 PFU/cell. Infections with NDV-GFP (Kindly provided by Dr. Adolfo García-Sastre, Icahn School of Medicine at Mount Sinai, New York, NY, United States) were performed at MOIs of 0.1 and 1PFU/cell.

### Antibodies and reagents

The following antibodies were used in Western blot: anti-β-actin mAb (Santa Cruz Biotechnology, #sc-47,778), rabbit polyclonal against the PKR protein (kindly provided by Dr. Javier Benavente, University of Santiago de Compostela, Spain), and a polyclonal serum against the IBDV VP3 protein (Fernandez-Arias et al., 1998).

Bafilomycin A1 (BFA) and staurosporine were purchased from Sigma-Aldrich, Poly I:C (high molecular weight) from InvivoGen, z-VAD-fmk (pan-caspase inhibitor) from Calbiochem, 7-deaza-2’-C-metiladenosina (7DMA) from Santa Cruz Biotechnology and ruxolitinib (Rx) from Selleckechem.

### Plasmids

The *chTLR3* gene fused to a His-tag sequence at the 3’end was chemically synthesized (GenScript) and cloned into the pcDNA3 vector (pc-chTLR3-His). The IFN-β reporter plasmid (pLUCTER) (King and Goodbourn, 1994) was kindly provided by Dr. Stephen Goodbourn. The *Renilla* luciferase plasmid (pRL-null; Promega) was kindly provided for Dr. Pablo Gastaminza. The NF-κB reporter plasmid (pSI-chNFκB-Luc) was generated as previously described [23].

### Virus titration

For extracellular virus titrations, media from infected cell cultures were collected at 16 and 24 h pi and were subjected to low-speed centrifugation (2,000xg for 5 min) to remove cell debris. The resulting supernatants were used to determine virus titers. For intracellular virus titrations cell monolayers were harvested by gentle scrapping, followed by low-speed centrifugation and pellets were suspended in fresh medium and subjected to three freeze and thaw cycles. Samples were subsequently centrifuged (2,000xg for 5 min) to dislodging virus particles from cell debris, and then used to determine intracellular virus titers. Virus titers were determined by plaque assay in QM7 cells using semisolid agar overlays followed by immunostaining as previously described [7]. For infections with NDV-GFP viral replication was estimated by measuring GFP expression. For this, fluorescence was quantified using a SpectraMax iD3 fluorometer (Molecular Devices). Cells were harvested as described above and cell pellets were resuspended in phosphate-buffered saline (PBS) and transferred to a black, clear-bottom 96-well microplate (Costar). Fluorescence was subsequently measured at an excitation wavelength of 395 nm and an emission wavelength of 509 nm.

### Luciferase reporter assays

Preconfluent (70-80% confluence) DF-1 cell monolayers were transfected in 24-well plates with either 100 ng of pLUCTER together with 30 ng of the pRL-null to normalize for transfection efficiency, or with 50ng of pSI-chNFκB-Luc, in combination with the pcDNA3-chTLR3-His or the empty pcDNA3 as control, as indicated in the results section, using Lipofectamine 2000 (Invitrogen) at a 1:2 ratio in Opti-MEM (Gibco) medium. 8 h later, cells were transfected with 250 ng of poly I:C. After 16 h, cells were lysed, and luciferase assays were performed with the dual-luciferase assay kit (Promega) according to the manufactureŕs instructions. Luciferase activity was recorded using an Appliskan luminometer (Thermo Scientific). Firefly luciferase values were normalized to *Renilla* values, and the fold induction was calculated as the ratio of samples transfected with poly I:C versus mock-transfected samples.

### Western blot assay

Infected cells were lysed in Laemmlís sample buffer (62.5 mM Tris-HCL [pH 6.8], 2% sodium dodecyl sulfate [SDS], 0.01% bromophenol blue, 10% glycerol, and 5% β-mercaptoethanol). Protein samples were subjected to 10% SDS-polyacrylamide gel electrophoresis (PAGE), followed by electroblotting onto nitrocellulose membranes (Bio-Rad). Membranes were incubated with blocking buffer (Tris-buffered saline [TBS] containing 0.05% Tween 20 [TBST] and 5% nonfat dry milk) for 30 min at room temperature, and incubated at 4°C overnight with the corresponding primary antibodies diluted in blocking buffer. Thereafter, membranes were washed with TBST and incubated with either a goat anti-rabbit IgG-peroxidase (Sigma) or a goat anti-mouse IgG-peroxidase conjugate (Sigma), and immunoreactive bands were detected by an enhanced chemiluminescence (ECL) reaction (SuperSignal Thermo Scientific) and recorded using the ChemicDoc Touch Imaging System (Bio-Rad).

### Caspase 3/7 activity assays

Quantification of caspase activity was carried out by using the Caspase-Glo 3/7 kit (Promega). For this, DF-1 cell monolayers grown in 24-well plates (70-80% confluence) were infected in duplicated. At 24 h pi, cells were harvested in medium and kept frozen until their analysis. 25 µl of the cell lysates under study was added to the same volume of Caspase-Glo 3/7 reagent in a 96-well plate. Plates were gently shaken and then incubated in the dark at room temperature for 1 h. The luciferase activity was recorded using an Appliskan luminometer (Thermo Scientific).

### Genome editing with CRISPR/Cas9

We used the Alt-R CRISPR-Cas9 system developed by Integrated DNA Technologies (IDT) to delete the two N-terminal dsRNA binding domains and a portion of the active site of the protein of PKR. Two sequence-specific CRISPR RNAs (crRNAs) for the *PKR* gene were designed using the Breaking-Cas website (https://bioinfogp.cnb.csic.es/tools/breakingcas/). The selected crRNA sequences were 1: CAAGCTCGATTACGTCGACGG and 2: TAGTGCGATATTATTGCAGCG. These crRNAs, the conserved transactivating crRNA (tracrRNA) required for the formation of the guide RNA (gRNA) and the Cas9 nuclease were purchased from IDT. Following the manufactureŕs protocol, tracrRNA and crRNA were combined, and resulting crRNA:tracrRNA duplexes were then mixed with the Cas9 to obtain ribonucleoprotein complexes (RNPs) that were transfected into the DF-1 cells plated in 6-well plates using Lipofectamine RNAiMAX (Invitrogen) in Opti-MEM medium according to the manufactureŕs instructions, and then seeded in 6-well plates. At 48 h pt, DF-1 cells monolayers were tripsinized, diluted and seeded in 100 mm dishes to obtain isolated clones. Individual clones were selected and grown, and subsequently analyzed by amplifying the regions of genomic DNA around the site of interest with the following primers: F1: 5’- GTGCATGGCCAAGATCAACAG-3’; R1: 5’-TCACGAGACCAAAGTCACCAA-3’ and R2: 5’- TGAGGAGGGCTAGTAAAACAGTAA-3’, used in two combinations, F1-R1 and F1-R2. Also, we analyzed the expression of PKR protein through Western blot. The results confirm the mutation introduced in the *PKR gene* and the abrogation of PKR expression.

In addition, we also generate DF-1 TLR3 KO cell lines. We used the same system to partially delete the extracellular domain of chicken TLR3. Three sequence-specific CRISPR RNAs (crRNAs) for the *chTLR3* gene were designed using the Breaking-Cas website. The selected crRNA sequences were 1: 5’-AACATGTTAGTGGTAATCCGTGG-3’; 2: GCCTAAATATCACGGTACTCTGG and 3: 5’-CAATTGCACGAACTC-CCTGATGG-3’. The following two pairs of crRNAs were used to generate different deletions in *TLR3* gene: crRNA 1 combined with crRNA 2, or crRNA2 together with crRNA 3. RNPs were obtained and transfected into cells as described above. The selected clones were analyzed by amplifying the genomic DNA with the following primers: F1: 5’- CAATAGTGGGGAAGCAGGAA-3’; F2: 5’-CCTTCCCGAATTTAGGCTTT-3’, and R1: 5’- TCCTGCTTCGAAGTCTCGTT-3’, used in two combinations, F1-R1 and F2-R1. The resulting PCR amplicons were sequenced to confirm the mutations introduced in the *TLR3* gene.

Following the same approach, we also generated DF-1 RIPK1 KO cell lines. Two sequence-specific crRNAs were designed to generate a large deletion encompassing from exon 2 to 12 in the *chRIPK1* gene, crRNA 1: 5’-GACAAACAACAATTAGATGCTGG-3’; crRNA 2: 5’- GATCACGACTACGAACGAGATGG-3’. Individual clones were selected and grown, and genetically characterized by amplifying the regions of genomic DNA around the site of interest with the following primers: F1: 5’-TATATGGCAGCGTCCCTCTC-3’, R2**: 5’-**TTGGCAGAAAAATCCATTCC-3’. The resulting PCR amplicons were sequenced to confirm the mutations introduced in the *RIPK1* gene.

### siRNA transfection

Silencing of endogenous *MDA5* and *MAVS* genes in DF-1 cells was performed using specific siRNAs, along with the respective non-targeting control siRNA (50 nM, Dharmacon^TM^). The cells in the 24-well plates were transfected with these siRNAs using Lipofectamine RNAiMax (Invitrogen), following the manufactureŕs instructions. The efficiency of silencing was checked by comparing mRNA expression levels of *MDA5* or *MAVS*, between silenced cells and control cells at different times pt by RT-qPCR The corresponding siRNAs sequences were 5’-AAGAAGGGAUCCAUUUAGA-3’ (siMDA5), 5’- UGCUGCAGGAAGCUUUGAA-3’ (siMAVS) [72], and 5’- AAGGACGCUGAGGCCUAAUCCUGUU-3’ (siControl) [73].

### RT-qPCR

Total RNA was isolated by using the NucleoSpin RNA plus (Macherey-Nagel) according to the manufactureŕs instructions. Purified RNA (250 ng) was reverse transcribed into cDNA by using random primers (ThermoFisher Scientific) and SuperScript III reverse transcriptase (Invitrogen), according to the manufactureŕs protocol. The cDNA was then subjected to RT-qPCR using the gene-specific primers indicated in **Table 1**. RT-qPCRs were performed in duplicate using Power SYBR green PCR master mix (ThermoFisher Scientific), according to the manufactureŕs protocol, on an Applied Biosystems 7500 real-time PCR system instrument. Reactions were performed as follows: 2 min at 50°C; 10 min at 95°C; 40 cycles of 15 s at 95°C and 1 min at 60°C; and finally, 15 s at 95°C, 1 min at 60°C, 30 s at 95°C, and 15 s at 60°C to build the melt curve. Gene expression levels were normalized to the *Glyceraldehyde-3-Phosphate Dehydrogenase (GAPDH)* gene, and the results were calculated as fold changes in gene expression relative to mock-infected cells by using the delta-delta *C_T_*(threshold cycle) method of analysis. Dilutions of plasmids containing the sequence amplified by each set of primers run in parallel were used to establish the corresponding standard curves.

**Table 1.**
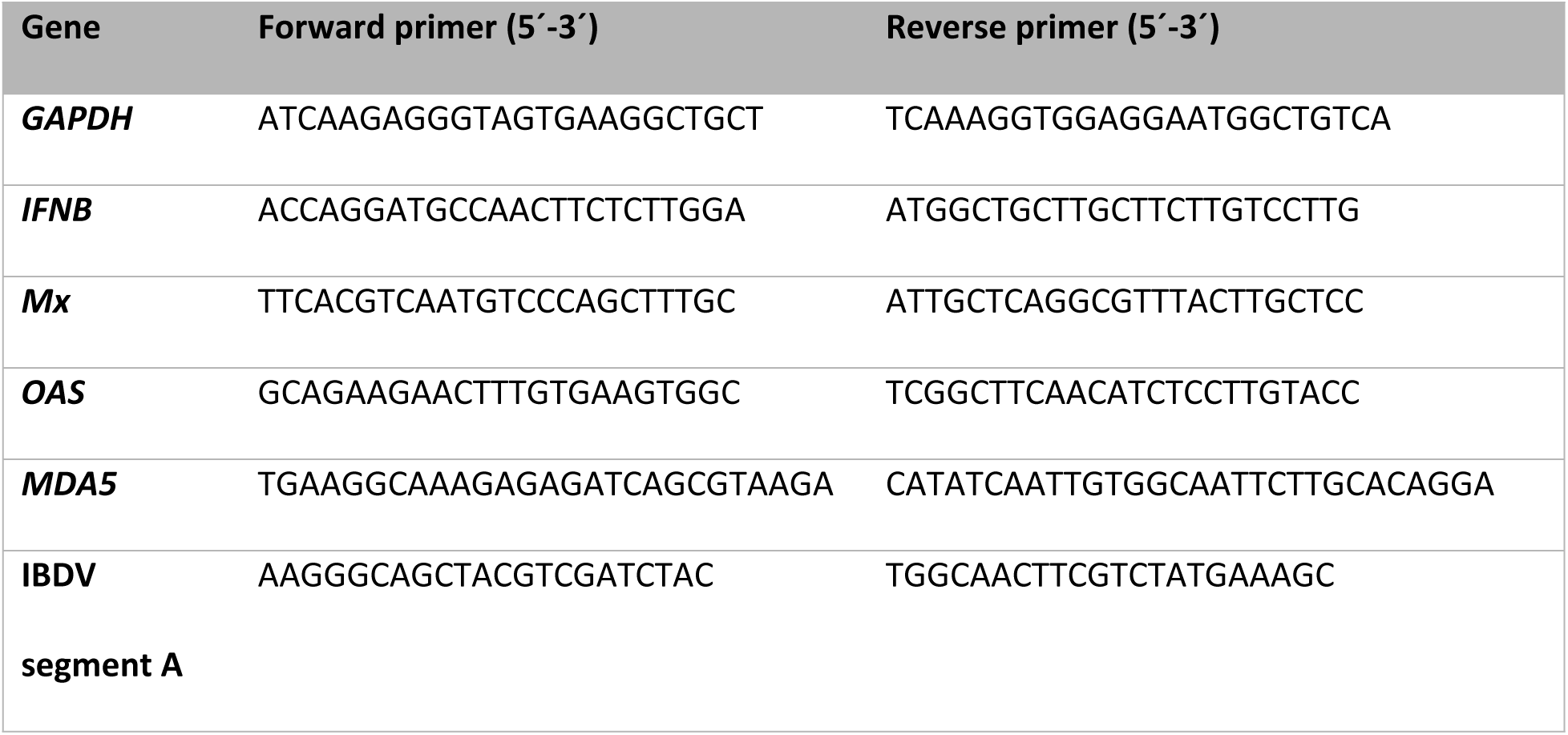
List of primers used for RT-qPCR.

### Statistics

GraphPad Prism version 5.03 software (GraphPad Software, La Jolla, CA) was used to determine statistical significance, using the Student unpaired two-tailed *t* test.

## Acknowledgements

We are grateful for the excellent technical assistance provided by Antonio Varas.

## Funding

This work was supported by Grants PID2020-112847RB-I00 to DR, JFR, JRR and FA, and CEX2023-001386-S to JFR, financed by MCIN/ AEI /10.13039/501100011033. ED-B was supported by the FPU 18/01873 research contract from the Spanish Ministry of Universities, AM-M was supported by Grant PRE2021-098502 funded by MICIU/AEI/10.13039/501100011033 and, as appropriate, by “ESF+”, and LC was recipient of contract 13-2022-008566 supported by the Community of Madrid through the Investigo Program, within the framework of the Recovery, Transformation and Resilience Plan, co-financed by the European Union through the NextGeneration EU funding initiative. The funders had no role in the study design, data collection and interpretation, or the decision to submit the work for publication.

---

**S1 Fig.** TLR3-dependent IBDV-induced cell death can be inhibited by the pan-caspase inhibitor Z-VAD-FMK. DF-1 and DF-1 TLR3 KO cells were mock-infected or infected with IBDV (MOI of 2 PFU/ml) in the presence or absence of the pan-caspase inhibitor Z-VAD-FMK (ZVAD) (50 µM). DF-1 and DF-1 TLR3 KO cells were also treated with staurosporine (STS) (1 µM) to confirm the ability of DF-1 TLR3 KO to undergo apoptosis in response to other stress inducing stimuli. Cells were harvested at 24 h pi and apoptosis was measured by using the Caspase-Glo 3/7 assay kit in duplicate. Caspase values from infected cell samples were normalized to those from mock-infected cells (Mock) (Striped bars). Bars indicate means ± standard deviations based on data of duplicate samples from three independent experiments. ** indicate *p* values of <0.01, as determined by unpaired Student’s test. ns, not significant.

**S2 Fig.** Analysis of the effect of TLR3 ablation on the production and release of IBDV VP5 KO compared to WT virus. DF-1 and DF-1 TLR3 KO cells were mock-infected or infected with IBDV WT or IBDV VP5 KO (MOI of 2 PFU/ml) and cells were harvested at 8 and 16 h pi for extracellular **(A)** and intracellular **(B)** virus titration. Bars indicate means ± standard deviations based on data of duplicate samples from three independent experiments. * and ** indicate *p* values of <0.05 and <0.01, respectively, as determined by unpaired Student’s test. ns, not significant.

## Notes

### Competing Interest Statement

The authors have declared no competing interest.

